# An atlas of genetic scores to predict multi-omic traits

**DOI:** 10.1101/2022.04.17.488593

**Authors:** Yu Xu, Scott C. Ritchie, Yujian Liang, Paul R. H. J. Timmers, Maik Pietzner, Loïc Lannelongue, Samuel A. Lambert, Usman A. Tahir, Sebastian May-Wilson, Åsa Johansson, Praveen Surendran, Artika P Nath, Elodie Persyn, James E. Peters, Clare Oliver-Williams, Shuliang Deng, Bram Prins, Carles Foguet, Jian’an Luan, Lorenzo Bomba, Nicole Soranzo, Emanuele Di Angelantonio, Nicola Pirastu, E Shyong Tai, Rob M van Dam, Emma E Davenport, Dirk S. Paul, Christopher Yau, Robert E. Gerszten, Anders Mälarstig, John Danesh, Xueling Sim, Claudia Langenberg, James F. Wilson, Adam S. Butterworth, Michael Inouye

## Abstract

Genetically predicted levels of multi-omic traits can uncover the molecular underpinnings of common phenotypes in a highly efficient manner. Here, we utilised a large cohort (INTERVAL; N=50,000 participants) with extensive multi-omic data for plasma proteomics (SomaScan, N=3,175; Olink, N=4,822), plasma metabolomics (Metabolon HD4, N=8,153), serum metabolomics (Nightingale, N=37,359), and whole blood Illumina RNA sequencing (N=4,136). We used machine learning to train genetic scores for 17,227 molecular traits, including 10,521 which reached Bonferroni-adjusted significance. We evaluated genetic score performances in external validation across European, Asian and African American ancestries, and assessed their longitudinal stability within diverse individuals. We demonstrated the utility of these multi-omic genetic scores by quantifying the genetic control of biological pathways and by generating a synthetic multi-omic dataset of UK Biobank to identify disease associations using a phenome-wide scan. Finally, we developed a portal (OmicsPred.org) to facilitate public access to all genetic scores and validation results as well as to serve as a platform for future extensions and enhancements of multi-omic genetic scores.

## Introduction

Multi-omic analysis has become a powerful approach to improve disease predictors and dissect the regulatory networks that underpin disease biology^1–3^. However, the collection of transcriptomic, proteomic, metabolomic and other modalities is an extremely expensive and time-consuming process. Because of these barriers, large-scale population cohorts typically generate multi-omic data for only a subset of participants (or not at all), which consequently reduces the statistical power of subsequent analyses and creates inequities for studies that do not have ample resources or are from underrepresented ancestries and other demographics.

It has been shown that genetic prediction of complex human traits can have both analytic validity and potential utility in research and clinical settings^4–8^. Genetic prediction has also been extended to omics data, for example whole blood^9^ and multi-tissue transcriptomics^10,11^ as well as plasma proteomics^12,13^. The value of such genetically-predicted traits is primarily in the elucidation of the molecular aetiology of common diseases, incorporating both directionality (as the germline genome is more or less fixed over a life course) and the power of large-scale genotyped biobanks to overcome prediction noise^14–16^.

The use of genetic scores to predict, expand and thereby democratize multi-omics data is an area of intense interest. While foundational, genetic prediction in this area has historically focused on gene expression, drawing on heterogeneous sources for training data which have limited sample sizes. With many cohorts now performing multi-omics profiling at scale, there is a unique opportunity to create genetic scores which capture multi-omic variation of population-based samples. Given suitably robust external validation, the reliability of multiomic genetic scores can be quantified and extended to analyses assessing their transferability across ancestries, thus facilitating equitable tools for molecular investigations in multiple populations. This approach both facilitates integrative cross-cohort analyses for multi-omic studies and enables the efficient generation of synthetic multi-omic data for studies with only genetic data assayed.

Here, we utilise the INTERVAL study^17^, a cohort of UK blood donors with extensive multiomic profiling, to train genetic prediction models. We externally validated these genetic scores in seven different external studies, comprising European, East Asian (Chinese, Malay), South Asian (Indian) and African American ancestries. We then demonstrate the use of genetically-predicted molecular data, including their coverage of biological pathways and the identification of multi-omic predictors of diseases and traits in UK Biobank. Finally, we construct an open resource (OmicsPred.org) which makes all genetic scores, validations and biomarker analyses freely available to the wider community.

## Results

### Development of genetic scores

This study aimed to develop genetic scores for blood biomolecular traits, including transcripts, proteins, metabolites (**Figure 1**). To do this, we used the INTERVAL study which collected participant serum or plasma on which assays from five different omics platforms were performed: SomaScan v3 (SomaLogic Inc., Boulder, Colorado, US), an aptamer-based multiplex protein assay; Olink Target (Olink Proteomics Inc., Uppsala, Sweden), an antibody-based proximity extension assay for proteins; Metabolon HD4 (Metabolon Inc., Durham, US), an untargeted mass spectrometry metabolomics platform; Nightingale (Nightingale Health Plc., Helsinki, Finland), a proton nuclear magnetic resonance (NMR) spectroscopy platform; and whole blood RNA sequencing via the Illumina NovaSeq 6000 (Illumina Inc., San Diego, California, US) (**Methods**). INTERVAL participants were genotyped on the Affymetrix Biobank Axiom array which was then imputed using a combined 1000 Genomes Phase 3-UK10K reference panel (**Methods**). After quality control, there were 10,572,788 genetic variants for constructing genetic scores.

**Figure 1:**
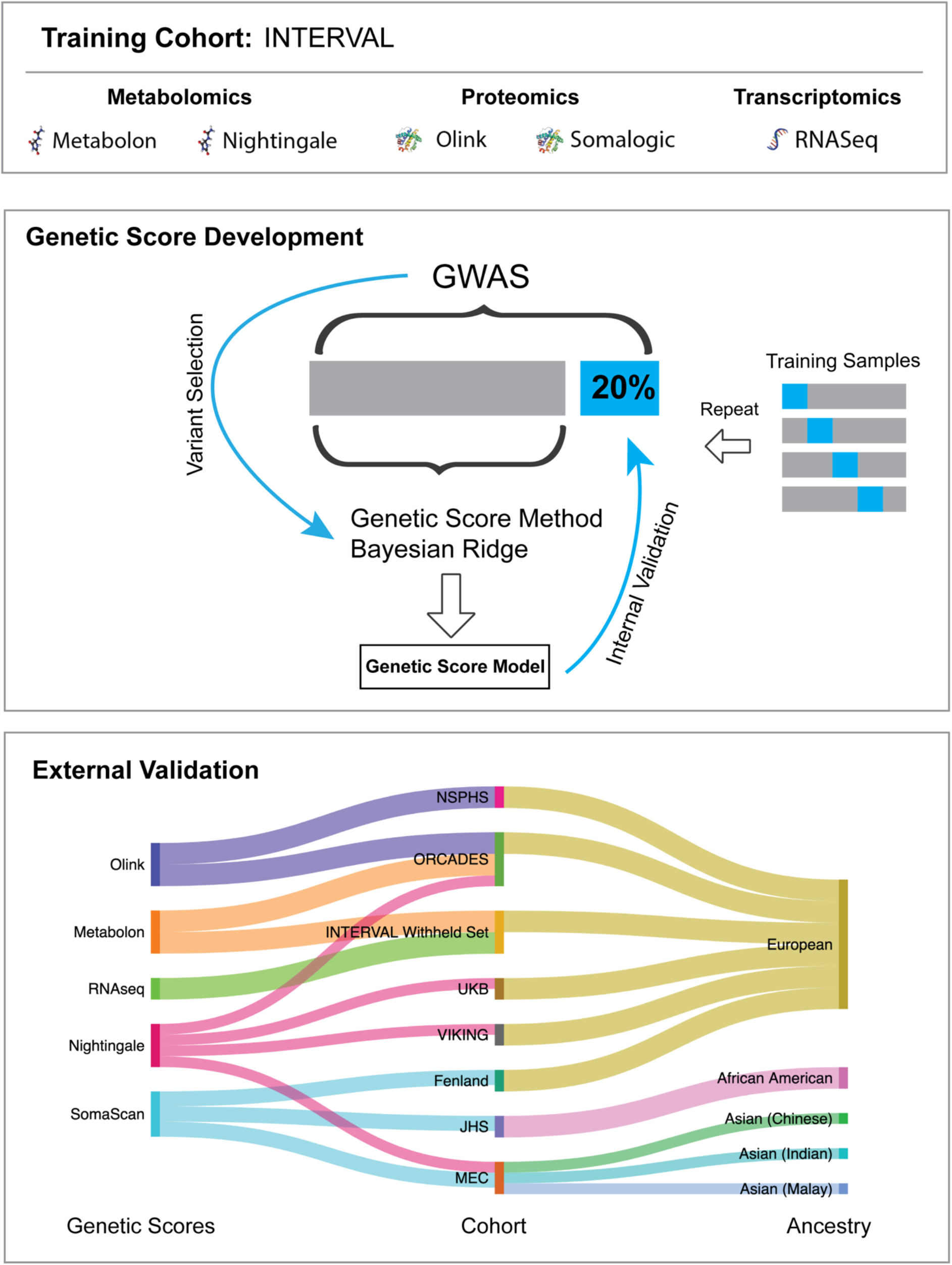
Schematic framework for the development and validation of multi-omic genetic scores.

To train genetic scores, we utilised Bayesian ridge regression (BR), which has been shown to have equal or better performance as other machine learning methods for genetic prediction^8^ and is more computationally efficient with a smaller carbon footprint^18^. In the data used here, we confirmed the generalisability of these findings across multiple platforms (Metabolon, Olink, SomaScan), assessing the impact of different sets of variants arising from different filtering strategies (**Methods**; **Figures S1-4**). Overall, we found the best performing approach overall to be BR with a genome-wide variant selection using GWAS p-value < 5×10^−8^ (**Figures S1-4**).

We developed genetic scores for 17,227 biomolecular traits from the five platforms, including 726 metabolites (Metabolon HD4), 141 metabolic traits (Nightingale), 308 proteins measured by Olink, 2,384 protein targets measured by SomaScan, 13,668 genes for Illumina RNAseq (Ensembl gene-level counts) (**Methods**). Across all platforms, we found wide variation in the predictive value (R^2^ between the genetically predicted and the directly measured biomolecular trait) and the number of variants of the genetic scores in internal validation (**Figure S5)**.

Overall, we found 10,521 biomolecular traits could be genetically predicted at Bonferroni-adjusted significance (correcting for all genetic scores tested), including 1,051, 206, 379, 137 and 8,748 for SomaScan, Olink, Metabolon, Nightingale and RNAseq respectively. Of these, 5,816 and 409 genetic scores could predict their biomolecular traits with *R*^*2*^ > 0.1 and *R*^*2*^ > 0.5, respectively (**Figure 2 and Tables S1-5**).

**Figure 2:**
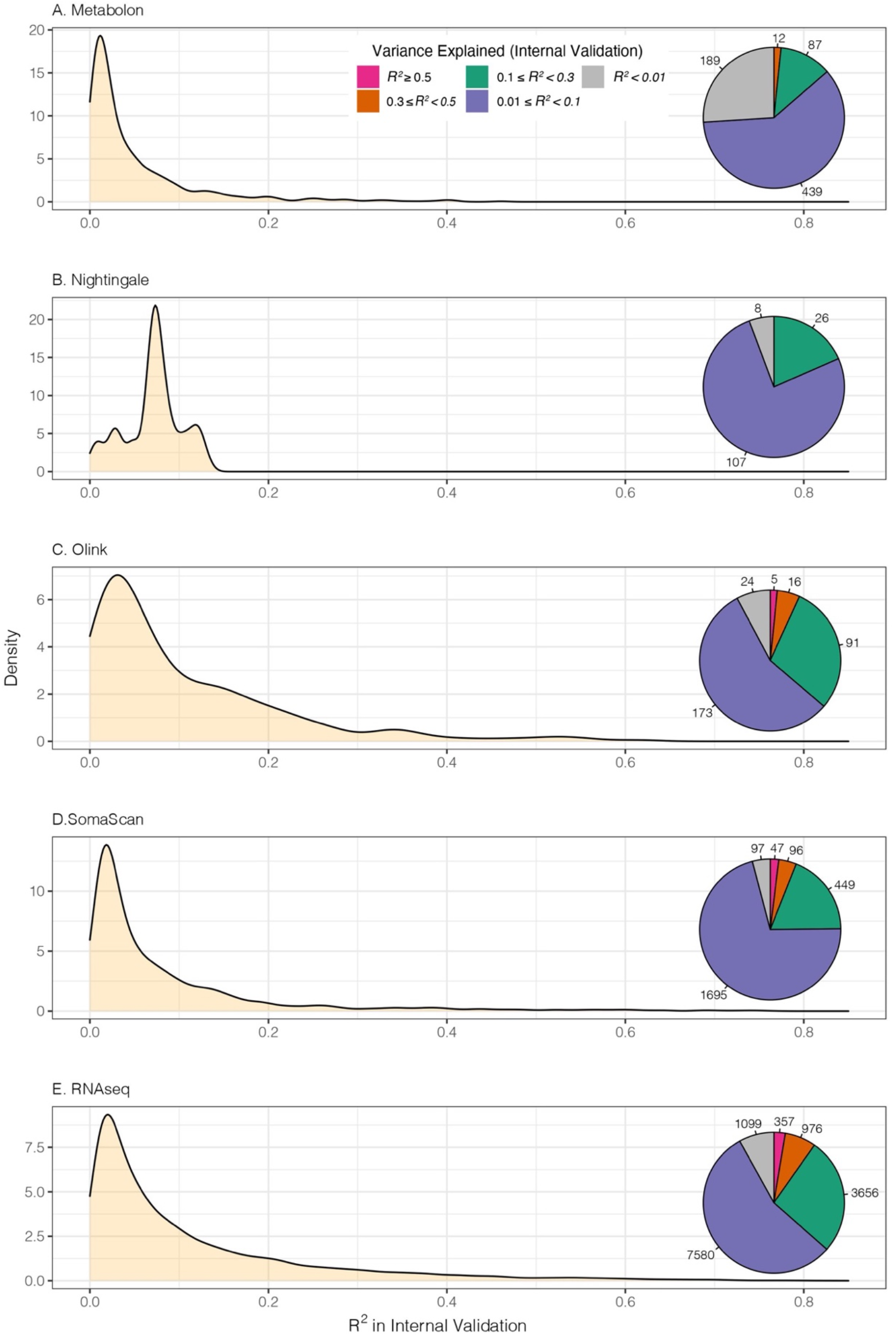
Performance of multi-omic genetic scores in internal validation. For each platform, genetic scores were constructed using Bayesian ridge regression on the genome-wide genetic variants with univariate p-value < 5×10^−8^ in INTERVAL. The variance explained in the measured biomolecular trait (R^2^) by the genetic score is assessed in the internal validation set (**Methods**). Pie charts reflect the number of genetic scores in a particular R^2^ range.

### Validation in external cohorts of European ancestries

Following internal validation of the genetic scores, we performed external validation of SomaScan protein targets in the FENLAND study^19^; Olink proteins in the Northern Swedish Population Health Study (NSPHS)^20,21^ and the Orkney Complex Disease Study (ORCADES)^22,23^; Metabolon metabolites in ORCADES^23^; Nightingale metabolic traits in UK Biobank (UKB)^24,25^, Viking Health Study Shetland (VIKING)^26^ and ORCADES^23^ studies (**Figure 1** and **Table 1**). For Metabolon metabolites and Illumina RNAseq transcripts, we performed further validation in withheld sets of INTERVAL (**Methods**). Overall, we found that performance of the genetic scores for most traits across the five platforms was consistent between internal and external validation in European ancestries, with genetic scores of many traits being highly predictive (**Figure 3** and **Figures S6-11**). As expected, we also found that genetic scores with high missingness rates amongst variants (e.g. due to allele frequency differences or technical factors) had attenuated power (**Methods; Figure S12**).

**Table 1.** Demographic statistics of training and validation samples for genetic score construction of blood biomolecular traits by platform. The table shows the mean ± standard deviation of age and BMI for participants in each cohort or cohort subset.

| Platform | Cohort | Ancestry | # Traits | # Samples | % Men | Age (years) | BMI (kg/m<sup>2</sup>) |
| --- | --- | --- | --- | --- | --- | --- | --- |
| Training and Internal Validation |  |  |  |  |  |  |  |
| Metabolon | INTERVAL | European | 726 | 8,153 | 51.0% | 43.9 ± 14.1 | 26.4 ± 4.6 |
| Nightingale |  |  | 141 | 37,359 | 51.0% | 43.7 ± 14.1 | 26.4 ± 4.6 |
| Olink |  |  | 308 | 4,822 | 59.3% | 59.0 ± 6.7 | 26.5 ± 4.1 |
| SomaScan |  |  | 2,384 | 3,175 | 50.8% | 43.6 ± 14.2 | 26.3 ± 4.7 |
| Illumina RNAseq |  |  | 13,668 | 4,136 | 56.4% | 54.6 ± 11.6 | 26.6 ± 4.4 |
| External Validation |  |  |  |  |  |  |  |
| Metabolon | INTERVAL withheld subset | European | 527 | 8,114 | 49.4% | 47.9 ± 13.8 | 26.5 ± 4.6 |
|  | ORCADES |  | 455 | 1,007 | 43.9% | 54.0 ± 15.3 | 27.7 ± 4.9 |
| Nightingale | UKB | European | 141 | 116,830 | 45.8% | 56.5 ± 8.1 | 27.4 ± 4.8 |
|  | ORCADES |  | 141 | 1,884 | 40.0% | 53.9 ± 15.0 | 27.8 ± 5.0 |
|  | VIKING |  | 141 | 2,046 | 39.9% | 49.8 ± 15.2 | 27.4 ± 4.9 |
|  | MEC | Chinese | 139 | 1,067 | 47.2% | 52.1 ± 9.9 | 23.5 ± 3.8 |
|  |  | Indian | 139 | 654 | 43.7% | 44.5 ± 11.6 | 26.4 ± 5.1 |
|  |  | Malay | 139 | 634 | 42.9% | 44.9 ± 11.1 | 26.9 ± 5.1 |
| Olink | NSPHS | European | 302 | 872 | 47.6% | 49.6 ± 20.2 | 26.7 ± 4.8 |
|  | ORCADES |  | 301 | 1,052 | 44.1% | 53.8 ± 15.7 | 27.7 ± 4.9 |
| SomaScan | FENLAND | European | 2,129 | 8,832 | 47.1% | 48.8 ± 7.4 | 26.9 ± 4.8 |
|  | MEC | Chinese | 2,070 | 645 | 46.0% | 51.9 ± 10.9 | 23.5 ± 3.9 |
|  |  | Indian | 2,070 | 564 | 45.0% | 44.0 ± 12.0 | 26.3 ± 5.3 |
|  |  | Malay | 2,070 | 563 | 43.9% | 44.4 ± 11.3 | 26.9 ± 5.2 |
|  | JHS | African American | 820 | 1,852 | 39.0% | 55.7 ± 12.8 | 31.6 ± 7.3 |
| Illumina RNAseq | INTERVAL withheld subset | European | 12,958 | 598 | 49.5% | 45.0 ± 13.1 | 26.8 ± 4.8 |

**Figure 3:**
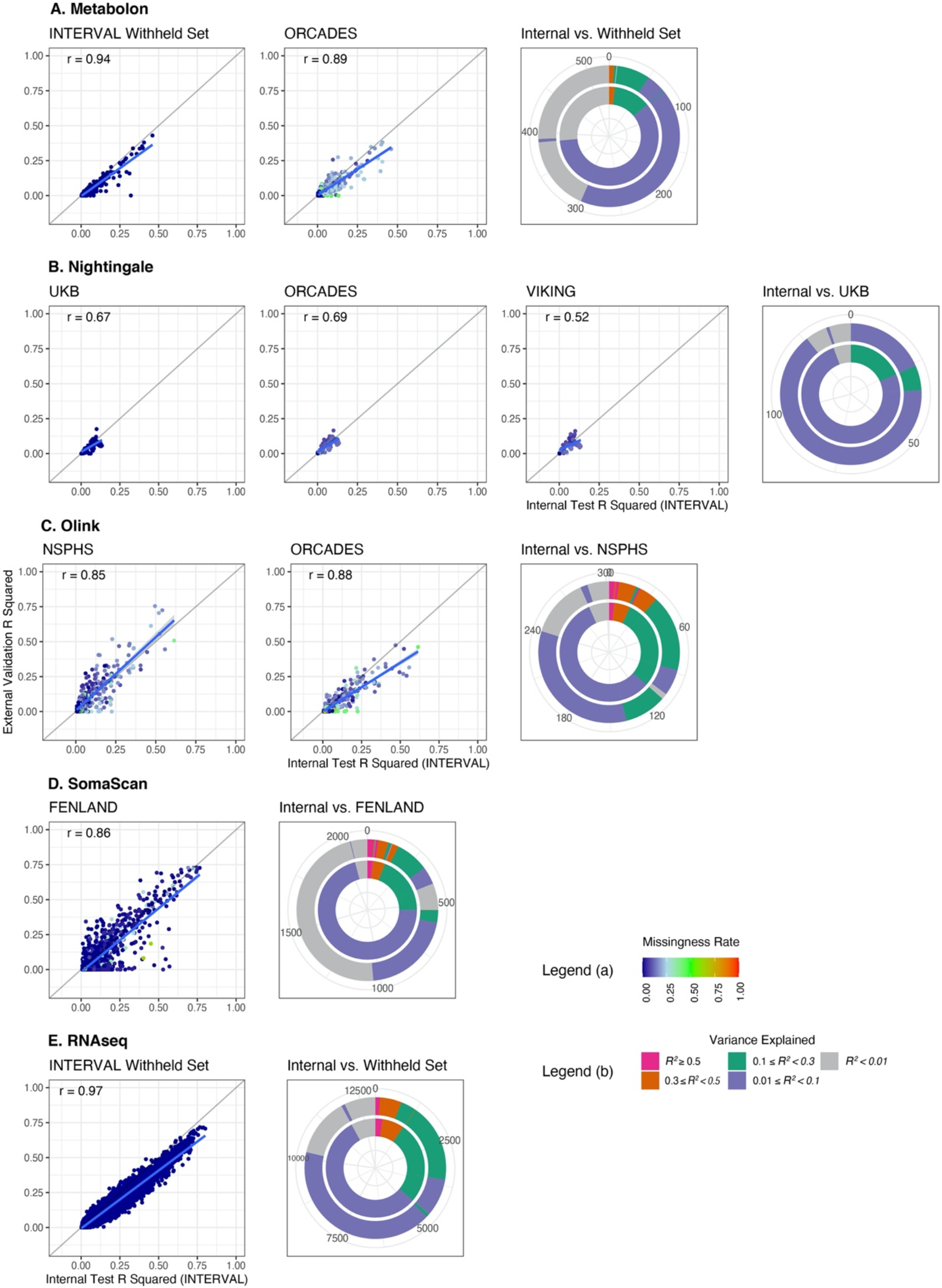
External validation of genetic scores in cohorts of European ancestry. Scatterplots show comparisons of the R^2^ in internal validation and external validation for each omic platform. Data points are coloured by variant missingness rate in the external cohort (i.e. the proportion of variants in the genetic score missing in the external cohort) and blue lines show the linear models fitting the data points. For each platform, concentric circles show the number of genetic scores in different ranges of explained variance (R^*2*^) in internal validation (inner ring) and external validation (outer ring). External validation cohorts used for each platform include FENLAND (SomaScan), NSPHS (Olink), INTERVAL withheld set (Metabolon), UKB (Nightingale) and INTERVAL withheld set (RNAseq).

The SomaScan v3 platform quantified 3,622 plasma protein targets in INTERVAL^27^, of which 2,384 proteins had at least one significant genetic variant that could be used for genetic score development (**Figure S5**). Internal validation found that SomaScan genetic scores had median R^2^ = 0.04 (IQR = 0.08). External validation in European ancestries utilised the FENLAND study^19^, where 89% (N=2,129) of SomaScan genetic scores could be tested. Overall, there was high consistency between internal and external R^2^ performance (Pearson correlation *r* = 0.86 across all SomaScan genetic scores tested) (**Figure 3**). Of the 2,129 tested SomaScan genetic scores, we found 45 proteins (2%) with a majority of their variance explained (R^2^ > 0.50) by the genetic score in external validation, including several involved in innate and adaptive immune responses, which were highly genetically predicted with R^2^ > 0.70 (CLEC12A, SIGLEC9, FCGR2A, FCGR2B and LILRB5). There were a total of 369 SomaScan proteins (17%) that could be genetically predicted with R^2^ > 0.10 in external validation.

The Olink proteomics used in INTERVAL quantified levels of 368 plasma proteins from four different panels (Inflammation, Cardiovascular 2, Cardiovascular 3, Neurology), of which 308 unique proteins were qualified for genetic score development (**Methods**). Internal validation found that Olink genetic scores had median R^2^ = 0.06 (IQR = 0.12). We were able to test 301 and 302 genetic scores in external European ancestry cohorts, NSPHS and ORCADES respectively (**Methods**). In assessing Olink proteins across both external validation cohorts, we found four proteins (FCGR2B, IL6R, MDGA1, SIRPA) with a majority of their variance explained (R^2^ > 0.50) by the genetic score in external validation, with FCGR2B on SomaScan found to be similarly genetically predicted (**Figure 3**). As compared to SomaScan, a larger proportion of Olink proteins in NSPHS (N=117; 39%) and ORCADES (N=87; 29%) could be genetically predicted with R^2^ > 0.10 in external validation. Overall, we found broad consistency between validations in NSPHS and ORCADES (**Figure S13**).

The Metabolon HD4 platform quantifies >900 plasma metabolites and was used here in two different phases of the INTERVAL study (**Methods**). Phase 1 (N=8,153) was used for development and internal validation of Metabolon genetic scores and phase 2 (N=8,114) was used for external validation (with no individuals overlapping between the two phases). We conducted a further external validation in ORCADES. Internal validation found that Metabolon genetic scores had median R^2^ = 0.02 (IQR = 0.05). A total of 726 Metabolon HD4 metabolites had significant genetic variants with which to construct genetic scores in INTERVAL, of which 526 and 455 metabolites (399 overlapping) could be externally validated in the phase 2 set and ORCADES, respectively (**Figure 3**). We again found broad consistency between the two external validation sets (**Figure S13**). There were no Metabolon HD4 metabolites with R^2^ > 0.50 between their genetic scores and their directly measured values in either the phase 2 set or ORCADES; however, there were 6 metabolites that had R^2^ > 0.3 in both the phase 2 set and ORCADES (4 metabolites overlapping). Of the metabolites that could be externally validated, 10% and 13% (N=50 and N=59) achieved a R^2^ > 0.10 in the phase 2 set and ORCADES, respectively. The top performing genetic scores included those for ethylmalonate (phase 2 set R^2^ = 0.43; ORCADES R^2^ = 0.33), N-acetylcitrulline (both phase 2 set and ORCADES R^2^ = 0.38) and androsterone sulfate (phase 2 set R^2^ = 0.35; ORCADES R^2^ = 0.17).

The Nightingale NMR platform was used to quantify 230 serum metabolic biomarkers (largely lipoproteins, lipids and low molecular weight metabolites) from 45,928 INTERVAL participants. Our analyses focused on the directly measured (non-derived) metabolic biomarkers, and genetic scores for 141 Nightingale biomarkers were developed using INTERVAL (**Methods**). Internal validation found that Nightingale genetic scores had median R^2^ = 0.07 (IQR = 0.03). The genetic scores were externally validated in three cohorts (UKB, ORCADES and VIKING). Overall, we found that genetic scores for Nightingale explained somewhat lesser variation in the directly measured traits, as compared to other platforms (**Figure 3**; **Figure S11**). Across UKB, ORCADES and VIKING, 28 Nightingale metabolic biomarkers had an R^2^ > 0.10 in at least one external validation cohort, with no biomarkers having R^2^ > 0.30. However, Nightingale genetic scores performed consistently across cohorts, with mean R^2^ for all 141 Nightingale biomarkers of 0.07, 0.06 and 0.06 in UKB, ORCADES and VIKING, respectively. The most predictive genetic scores were mainly related to low-density lipoprotein (LDL), e.g. concentrations of cholesteryl esters in small LDL, cholesterol in small LDL, cholesteryl esters in medium LDL, cholesterol in medium LDL and LDL cholesterol (**Table S2)**.

RNAseq of whole blood from 4,778 individuals in INTERVAL was carried out using Illumina NovaSeq (**Methods**). While 4,136 individuals were used to develop and test genetic scores, 598 individuals were kept as a withheld set for validation. The INTERVAL RNAseq data allowed for the construction of genetic scores using both *cis* and *trans* eQTLs for 13,668 genes (ENSEMBL gene IDs), of which 12,958 (95%) could be assessed in the withheld validation set (**Figure 3**). Internal validation found that RNAseq genetic scores had median R^2^ = 0.06 (IQR = 0.13). Overall, we found strong correlation of R^2^ between the internal and withheld validation sets (Pearson r = 0.97). There were 141 genes which had R^2^ > 0.50 in the withheld validation set, and 798 genes with R^2^ > 0.30. The most predictive genes were those involved in proteolysis (*RNPEP*; R^2^ = 0.71), solute cotransport (*SLC12A7*; R^2^ = 0.72), RNA helicase activity (*DDX11*; R^2^ = 0.71) and spliceosome function (*U2AF1*; R^2^ = 0.72).

### Transferability of multi-omic genetic scores to African American and Asian ancestries

To assess the performance of the genetic scores developed in the predominantly-European INTERVAL cohort in non-European ancestries, we utilised the Singapore Multi-Ethnic Cohort (MEC)^28^ and the Jackson Heart Study (JHS)^29^. MEC data comprised individuals of Chinese, Indian and Malay populations who have matched genotypes, plasma Nightingale NMR and plasma SomaScan (**Table 1; Methods**). The JHS data comprised African Americans with matched genotypes and plasma SomaScan (**Table 1; Methods**).

Overall, we found that genetic scores developed from INTERVAL can predict the Nightingale and SomaScan trait levels in cohorts of Asian and African American ancestries, but as expected their performances were significantly reduced when compared to the validations in European ancestry cohorts (**Figure 4**). For Nightingale, the European-trained genetic score performance generally declined from Chinese to Indian to Malay ancestries, with LDL subclasses displaying some of the most variable cross-ancestry R^2^ (**Figure 4a** and **4b**). The most transferrable Nightingale genetic scores were triglycerides in IDL, triglycerides in small HDL and medium HDL, degree of unsaturation and phosphatidylcholines (**Figure 4c**). When assessing transferability of SomaScan, we found genetic score performance generally declined from Indian to Malay to Chinese to African American ancestries (**Figure 4d**). The SomaScan genetic scores that attenuated most in non-European ancestries were those for CD177 (a cell-surface expressed protein on neutrophil and Treg’s) and GDF5 (a secreted ligand of TGF-beta) (**Figure 4e**). The most transferable SomaScan genetic scores included SIGLEC9 (which mediates sialic-acid binding to cells), SIRPA (a cell surface receptor for CD47 involved in signal transduction) and ACP1 (an acid and protein tyrosine phosphatase), where all internal and external validation R^2^ were >0.50 (**Figure 4f**).

**Figure 4:**
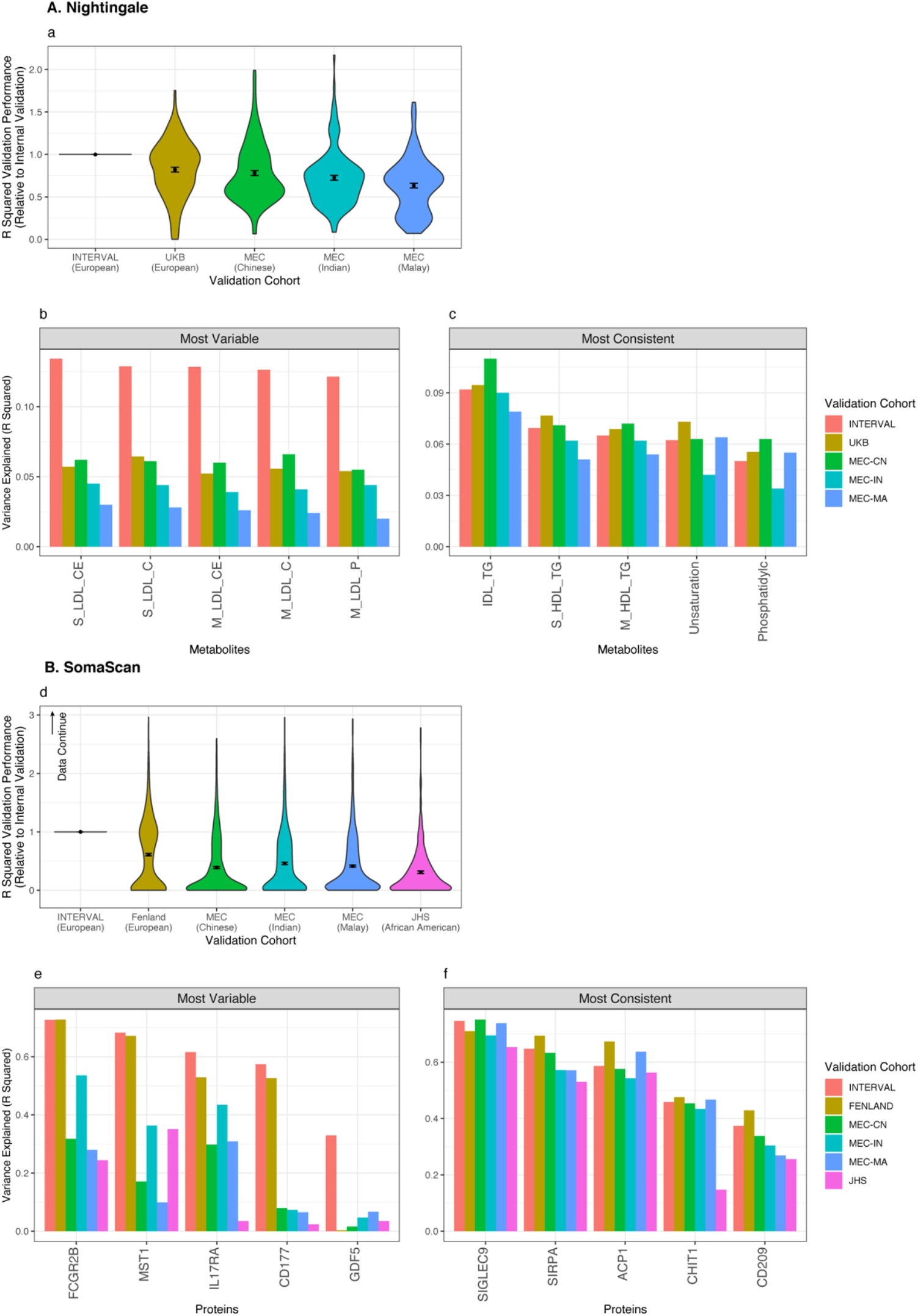
Transferability of genetic scores to cohorts of Asian and African American ancestries. (**a, d**) Performance (R^2^) of genetic scores for Nightingale (**a**) and SomaScan (**d**) in external cohorts of various ancestries relative to R^2^ in internal validation (INTERVAL). Transferability was only tested if the genetic score had a significant (Bonferroni corrected p-value < 0.05) association with the directly measured molecular trait in internal validation, which resulted in 137, 136 Nightingale metabolic traits for UKB and MEC (Chinese, Indian and Malay) respectively and 949, 945, 378 SomaScan proteins for FENLAND, MEC and JHS. Violin plots show distributions of the ratio of R^2^ values. Black points show mean values and error bars are standard errors. (**b, c, e, f**) R^2^ of genetic scores for Nightingale (**b, c**) and SomaScan (**e, f**) with the five most variable (**b, e**) or five most consistent (**c, f**) for prediction in multi-ancestry validation, as quantified by mean absolute difference in R^2^. In this analysis, only Nightingale R^2^ > 0.05, SomaScan R^2^ > 0.30 in internal validation were considered.

### Longitudinal stability of genetic scores in diverse ancestries

Within MEC, 1,739 individuals were measured at both baseline and revisit with mean length of follow-up 6.31 years (SD 1.45 years). This allowed longitudinal assessment of the stability of genetic scores for SomaScan (N = 403 Chinese, 356 Indian and 353 Malay) and Nightingale (N = 721 Chinese, 376 Indian and 363 Malay) platforms. For SomaScan traits, we found strong consistency between the predictive capacity of genetic scores between baseline and revisit samples (Pearson r = 0.99 for Chinese, 0.98 for Indian and 0.98 for Malay populations), and little difference in longitudinal stability between ancestries (**Figure 5d-f**). For Nightingale traits, despite variation in the predictive capacity of genetic scores between baseline and revisit samples, the longitudinal stability between ancestries was still largely consistent (Pearson r = 0.60 for Chinese, 0.84 for Indian and 0.85 for Malay populations; **Figure 5a-c**).

**Figure 5:**
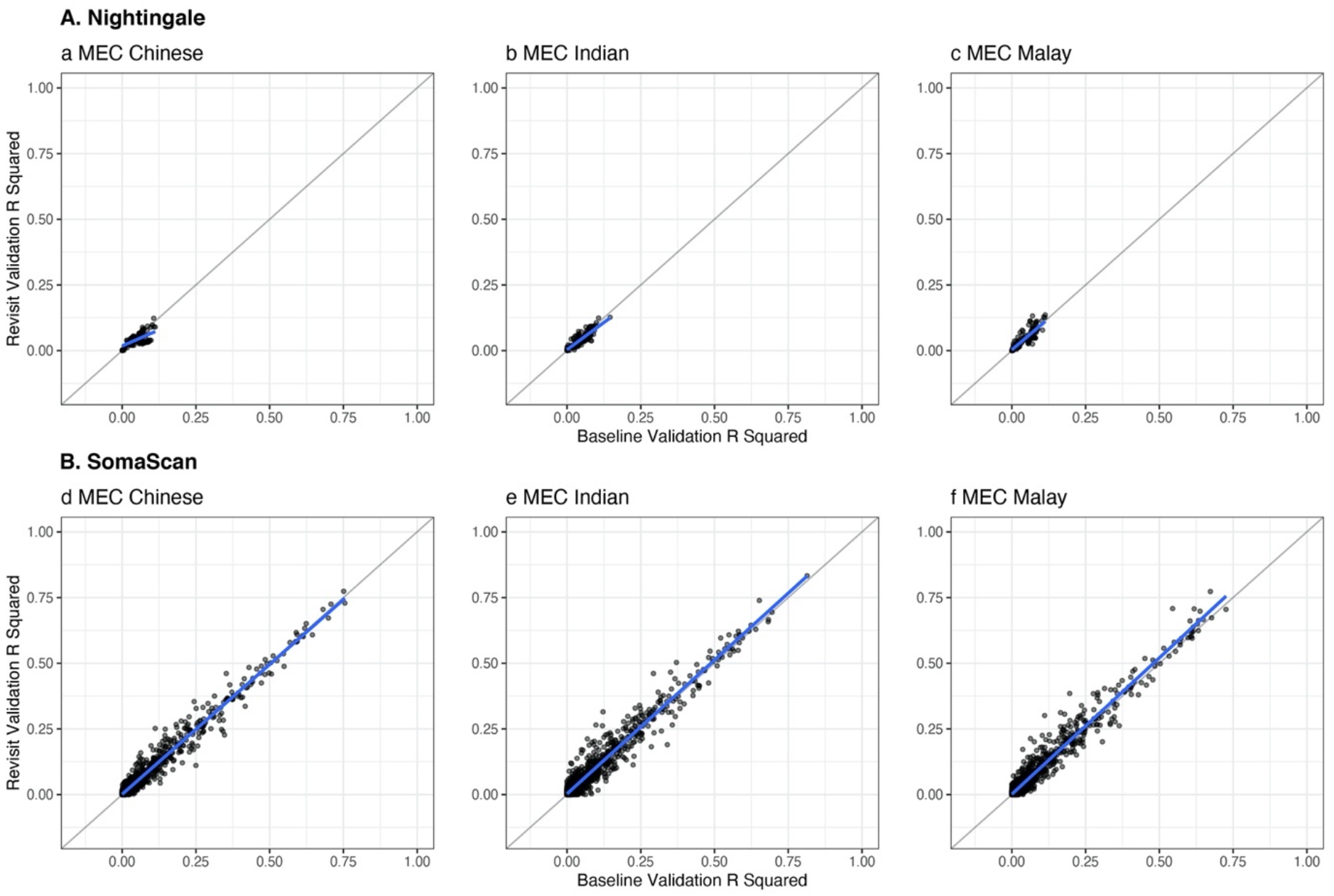
Performance (R^2^) of genetic scores between longitudinal samples and across ancestries in the MEC cohort. Paired samples include a baseline and a revisit sample from each individual run on Nightingale and SomaScan for MEC Chinese (N=406 and 721 individuals), MEC Indian (N= 356 and 376) and MEC Malay (N=353 and 363). Blue lines denote linear models fitted to each set of data points.

### Quantifying the genetic control of biological pathways

Multi-omic genetic scores may be used to probe the relevance of biological pathways to a particular trait or disease outcome of interest. To assess the coverage of biological pathways by the proteomic genetic scores we present here, we applied the genetic scores for SomaScan and Olink to assess the extent to which pathways are genetically controlled (**Methods**). Here, we considered all genetic scores with R^2^ > 0.01 in internal validation (2,205 unique proteins) and jointly mapped the SomaScan and Olink scores onto data curated from Reactome^30^ (**Figure 6a, Figure S15**).

**Figure 6:**
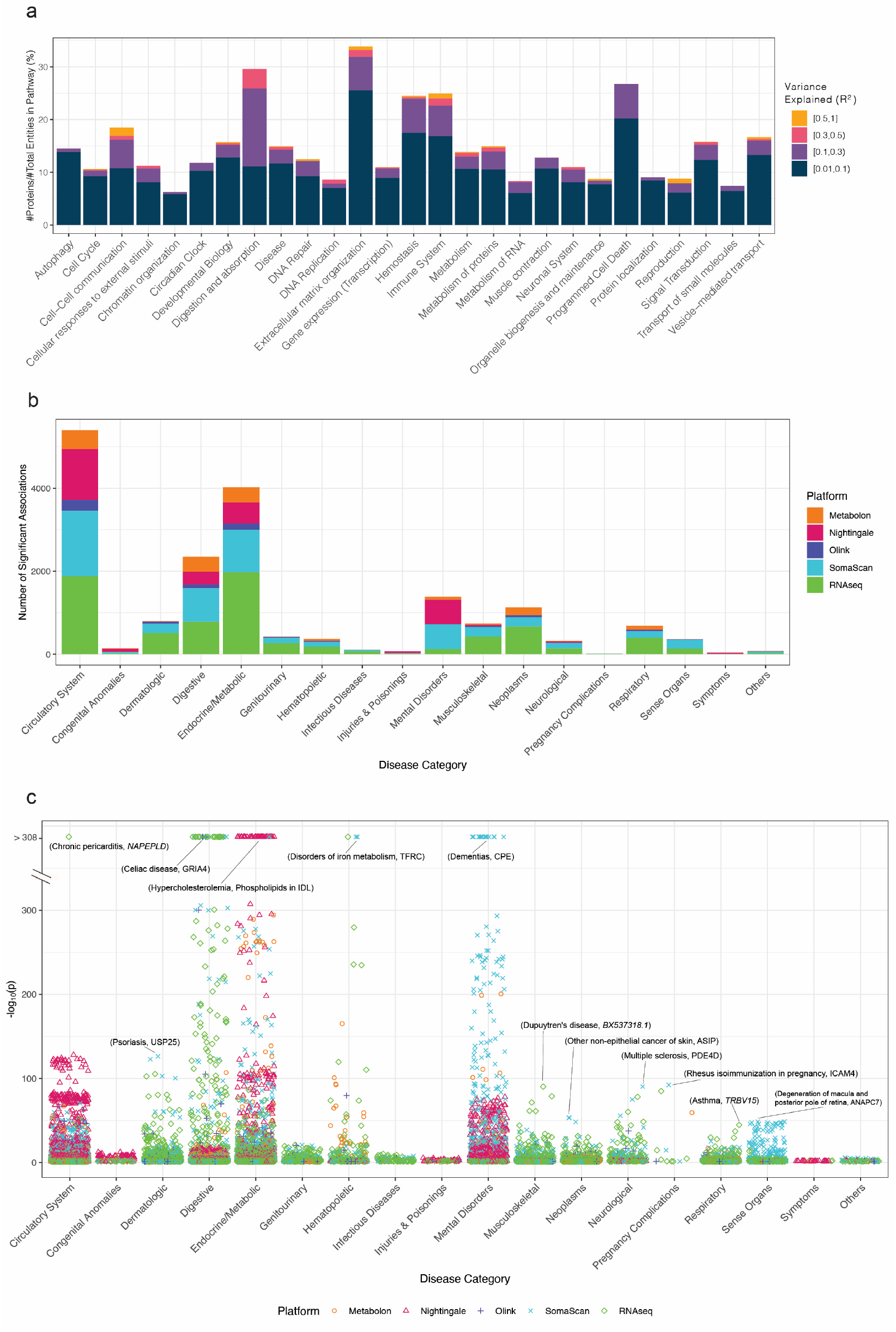
Applications of multi-omic genetic scores. (a) Genetic control of Reactome super-pathways using SomaScan and Olink genetic scores of varying predictive performances in internal validation (**Methods**). (b) Phenome-wide association study using PheCodes in UK Biobank. Slacked barplots showing the number of detected significant associations (FDR-corrected p-value < 0.05) by PheCode category of disease and omic platform. (c) Strength of associations by category of disease and omic platform. Association with the lowest pvalue for each category of diseases is labelled.

For the plasma proteome, we found wide variation amongst the 27 super-pathways with some super-pathways under relatively little genetic control (e.g. chromatic organisation, or transport of small molecules) and others under substantially greater genetic control (e.g. digestion and absorption, or extracellular matrix organisation) (**Figure 6a**). Approximately 18% of proteins in the digestion and absorption super-pathway had internal validation R^2^ > 0.10, and ∼4% with R^2^ > 0.30. For the lowest-level pathway annotation (N=1,717) of the 27 super-pathways, we found that a majority (N=1,169, 68%) were covered by at least one SomaScan or Olink genetic score with an internal validation R^2^ > 0.01 (**Figure S15)**. For both the digestion and absorption and the extracellular matrix organisation super-pathways, 25% and 42%, respectively, of lowest-level pathway annotations were covered by at least one SomaScan or Olink genetic score with internal R^2^ > 0.30.

### Phenome-wide association analysis using multi-omic genetic scores

Using the multi-omic genetic scores, we generated genetically predicted Metabolon HD4, Nightingale NMR, Olink, SomaScan and whole blood RNAseq data for the UK Biobank (**Methods**). Next, using these predicted multi-omics data of UKB, we performed a phenome-wide association study using PheCodes^31^ (ICD-9 and ICD-10 based diagnosis codes collapsed into hierarchical clinical disease groups; **Methods**). For simplicity and to maximize the number of qualified PheCodes, we focused the analysis on UKB individuals of white British ancestry. Multiple testing was controlled using Benjamini-Hochberg FDR of 0.05 (**Methods**).

Overall, at an FDR of 5%, we identified 18,404 associations between genetic scores of the biomolecular traits and 18 categories of PheCodes (**Figure 6b**). These associations comprised 1,668 for Metabolon HD4, 2,854 for Nightingale NMR, 740 for Olink, 5,501 for SomaScan and 7,641 for RNAseq (**Table S6** and **S7**). Circulatory system diseases, endocrine/metabolic and digestive diseases yielded the largest number of associations across platforms (**Figure 6b**).

The PheWAS detected many well-known blood biomarkers as well as intriguing associations across genes, proteins and metabolites. For example, total cholesterol was significantly associated with myocardial infarction (HR = 1.13 per s.d., FDR-corrected p-value = 1×10^−61^). Interleukin-6 (IL-6) pathways have been shown to have a causal association with coronary artery disease^32,33^, and notably, IL-6 receptor genetic scores in SomaScan and Olink had R^*2*^ > 0.50 in both internal and external validation, showing its high genetic predictability. Genetically predicted levels of IL-6 receptor in both Olink and SomaScan were significantly associated with myocardial infarction (HR = 0.97 per s.d., FDR-corrected p-value = 2×10^−4^; HR = 0.97 per s.d., FDR-corrected p-value = 4×10^−4^, respectively). Microseminoprotein-beta has been identified as a biomarker for prostate cancer^34^ and PheWAS findings support this association (HR = 0.87 per s.d., FDR-corrected p-value = 3×10^−49^). Genetically predicted Sex Hormone-Binding Globulin (SHBG) protein was associated with type 2 diabetes (HR = 0.98 per s.d., FDR-corrected p-value = 0.03), consistent with previous observational and genetic analyses^35^. Similarly, we found associations for insulin signaling pathway related proteins, e.g. insulin receptor (INSR) and insulin-like growth factor 1 receptor (IGF1R), with type 2 diabetes^36,37^; ABO^38^ with type 2 diabetes; IL-6 with asthma^39^; and *HLA-DQA1*/*DQB1* with celiac disease^40^ (**Table S6**).

Our results validate those of a recent study identifying putative causal plasma protein mediators between polygenic risk and incident cardiometabolic disease^5^, including six of the novel and putatively causal associations for coronary artery disease (**Table S6**). Amongst the strongest signals, we found intriguing associations including chronic pericarditis (N=266 cases) with genetically-predicted gene expression of the phospholipase *NAPEPLD* (HR = 0.88 per s.d., FDR-corrected p-value < 1×10^−307^) and the association of rhesus isoimmunization in pregnancy (i.e. maternal antibodies attacking fetal blood cells; N=302 cases) with genetically-predicted protein levels of ICAM4 (HR = 0.19 per s.d., FDR-corrected p-value = 3×10^−93^). ICAM4 itself is critical to the Landsteiner-Weiner blood system, which is genetically independent of the rhesus factor (Rh) blood group system. Despite the *ICAM4* locus showing no significant association with rhesus isoimmunization in pregnancy (PheWeb^41^), our ICAM4 results demonstrate that genetic prediction of plasma protein levels can identify biologically plausible candidate associations.

### OmicsPred: An online portal for multi-omic genetic scores

We developed an online portal (OmicsPred.org) to facilitate open dissemination of the genetic scores, detailed validation results and visualisations. OmicsPred also serves as an online updatable resource, which allows future expansion and deepening of the omics platforms, multi-ancestry transferability, newly developed and more powerful genetic scores, as well as results from applications of OmicsPred (**Figure S14)**.

The portal presents genetic scores of biomolecular traits by platform, in which users can access summary statistics of the training and validation cohorts used for traits at each platform as well as download, individually or in batch, the corresponding model files for genetic scores (i.e. variants and weights). Users can visualise validation results by selected performance metrics (e.g. R^2^ or Spearman’s rho), cohort(s), together with detailed trait (e.g. full protein name) and validation information (e.g. variant missingness rate). Users can easily search the portal to find biomolecular traits of specific interest, either by name or related descriptions. The OmicsPred portal also hosts descriptions and summary results from applications of the genetic scores (e.g. the PheWAS in UK Biobank described above).

## Discussion

In this study, we developed genetic scores for >17,000 multi-omic traits across five molecular platforms covering proteomics, metabolomics and transcriptomics in a single cohort. The relative predictive values and robustness of the genetic scores were assessed in external validations of European, Asian and African American ancestries; the longitudinal stabilities of the genetic score performances were established within individuals of different ancestries; and the utility of the multi-omic genetic scores was demonstrated by elucidating the relative genetic control of biological pathways and by identifying multi-omic disease associations using a phenome-wide scan of predicted multi-omic data in UK Biobank. Finally, we developed an open resource OmicsPred (OmicsPred.org) to publicly disseminate and continuously enhance the value of multi-omic genetic scores.

While the utility of generating predicted transcriptomic data for cohorts with genome-wide genotype data has been demonstrated^42^, our work substantially extends these foundations using a large multi-omic cohort, quantifying both the intra-and inter-ancestry reliability of proteomic and metabolomic genetic scores across multiple platforms. We generate a predicted multi-omic dataset for UK Biobank and show that PheWAS can uncover many known and novel omic associations with disease. Given that the increase in sample size required to detect an association for a noisy explanatory variable can be estimated by the formula n/R (where n is the sample size required if no measurement error exists and R is the reliability coefficient)^14^, even genetic scores of apparently low predictive value are likely powerful enough to detect true associations at the sample sizes of current and forthcoming biobank-scale data. This suggests that large biobanks could reliably test trait-disease associations using efficient genetically-predicted data, before committing to novel data generation using (frequently expensive) molecular assays.

Our study has several limitations. While blood is a key tissue of broad utility in discovery science and medicine, it is most likely a correlate but not the main site of function for many of the biomolecules assessed here. While genetic score performance was generally consistent across cohorts, there were factors that could affect their performance, including technical factors (e.g. use of serum versus plasma; genetic variant missingness), participant demographics, and genetic factors (e.g. allele frequency differences). Genetic scores may also pick up differences in molecular traits shared by multiple platforms (e.g. Olink and SomaScan). Despite genetic scores for most shared proteins being consistently predictive across platforms, there can be large differences which can be due to technical factors (e.g. binding affinity) (**Methods**), as assessed in a recent study^43^. The attenuated performance of polygenic scores across ancestries is a well-known limitation^44^ and our analysis also found this in multi-omics data. Multi-omics for non-European ancestries will likely become more common in the future, and we see a key role for OmicsPred in facilitating robust genetic scores which enable multi-omic prediction in diverse populations. Finally, we acknowledge that there are many highly sophisticated machine learning approaches, which may improve genetic score performance and/or transferability. We selected Bayesian ridge because it has been previously shown to both perform well relative to other machine learning approaches and because it scales very well to large numbers of traits, thus improving computational efficiency and promoting green computing^8,18,45^. Optimal variant selection thresholds may also vary for each platform or trait and this could potentially led to some improvements in prediction.

Future avenues for research include assessing to what extent the predicted multi-omic associations are causal, expansion of OmicsPred to additional platforms and/or cohorts, and multi-ancestry training for improved prediction. In summary, we have developed, validated and applied multi-omic genetic scores for >17,000 traits and made them publicly accessible via the new OmicsPred resource (https://www.omicspred.org), facilitating the generation and application of multi-omics data at scale for the wider community.

## Methods

### INTERVAL cohorts and data quality control

The INTERVAL study^17^ is a randomised trial of ∼50,000 healthy blood donors, who were recruited at 25 centres of England’s National Health Service Blood and Transplant (NHSBT) and aged 18 years or older at recruitment. This trial aimed to study the safety of varying frequency of blood donation, and all the participants completed an online questionnaire when joining the study about their demographic and lifestyle, such as age, sex, weight, height, alcohol intake, smoking habits, and diet, etc. All participants have given informed consent and this study was approved by the National Research Ethics Service (11/EE/0538).

Using the aptamer-based SomaScan assay (version 3), this study profiled plasma proteins of 3,562 participants in two batches (n=2,731 and n=831), of which 3,175 samples remained for analysis after quality control. The detailed steps for measurements and quality controls of the protein levels using the SomaScan array in INTERVAL have been previously described^5,27^. In summary, the relative concentration of 3,622 proteins (or protein complexes) targeted by 4,034 modified aptamers (*SOMAmer reagents*, referred to as SOMAmers) on the array were measured from 150-μl aliquots of plasma at SomaLogic Inc. (Boulder Colorado, US). Quality control was performed at the sample and SOMAmer levels by Somalogic, which uses the control aptamers and calibrator samples to correct for systematic variability in hybridization, within-run and between-run technical variability. For this study, we did not exclude protein aptamers with greater than 20% coefficient of variation in either batch, but excluded these aptamers targeting non-human proteins. We also excluded aptamers that, since the original quantification in INTERVAL, had been (1) deprecated by SomaLogic; (2) found to be measuring the fusion construct rather than the target protein; or (3) measuring a common contaminant^5^, which finally filtered the data to 3,793 high quality aptamers targeting 3,442 proteins. Within each batch, the relative protein abundances were natural log-transformed, and then adjusted for age, sex, the first three genetic principal components and duration between blood draw and sample processing (binary, 1 day vs >1 day). The protein residuals from this linear regression were finally rank-inverse normalized and used as phenotype values for their GWAS, which has been previously reported in detail^27^. These normalized phenotype values were further adjusted for batch effect and top 4-10 genetic principal components, which were used as the phenotype values for the genetic score model training and internal validation.

Using Olink proximity extension assays^46^, the INTERNAL study measured plasma protein abundance of ∼5,000 samples on four Olink panels: *Inflammation-1* (INF-1), *Cardiovascular II* (CVD-2), *Cardiovascular III* (CVD-3), and *Neurology* (NEUR) panel, each of which includes 92 proteins. For the INF-1, CVD-2 and CVD-3 panels, samples were assayed in two equal batches and their protein levels were pre-processed and quality controlled by Olink using NPX Manager software. Protein levels were then regressed on age, sex, sample measurement plate, time from blood draw to sample processing (number of days), season (categorical: spring, summer, autumn, winter), and inverse rank normal transformed. Details of quality control and GWAS for proteins on these three panels were given in the previous work^13^. Due to timing and funding differences, the NEUR panel was treated separately from other 3 panels for QC purposes. In detail, samples were assayed in one large batch, and trait levels were also processed by the NPX software and final measurements were presented as NPX values on a log^2^ scale (i.e. a one unit increase represents a doubling of protein level). We removed 187 measurements flagged by Olink as potentially having technical issues and 147 samples of potentially non-European origin as determined by principal component analyses, which left 4,811 measurements proceeding to standard QC assessments. We also checked for missing measurements and measurements below the limit of detection. No missing measurements were found. 8 out of 92 proteins had values below the limit of detection (LOD), of which 4 (HAGH, BDNF, GDNF, CSF3) had more than 5% of measurements below the LOD so were not taken forward for further analyses. No participant had more than 4% of protein measurements below LOD, and we did not observe over-representation of particular proteins below LOD for specific participants. Protein measurements were then adjusted for age, sex, season and the first 11 genetic PCs, residuals of which were further inverse normal rank transformed for their GWASs. It was noted that there are a small number of shared proteins across the four Olink panels (detailed numbers of proteins and participants per panel after QC were given in **Table S8**). To avoid duplication in genetic score construction, these shared proteins were merged by averaging their protein levels on each sample across panels, and taken as a unique protein. All the genetic variants identified in GWASs for the same protein across multiple panels were combined (if different) for its genetic score development. The normalized proteins levels of 308 unique proteins were adjusted for the first ten genetic principal components (if not adjusted previously), which were used as phenotype values for genetic score model construction and testing in INTERVAL.

The DiscoveryHD4® platform (Metabolon, Inc., Durham, USA) was used to measure plasma metabolites of INTERVAL participants. Four sub cohorts of 4,316 4,637, 3,333 and 4,802 participants were created through random sampling from the INTERVAL study and metabolites were measured within the four sub cohorts (or batches) separately at two time phases of the study (two batches at each phase). Samples of the first two batches were used as training data for GWAS and genetic score development of metabolite traits in the platform, and samples of the other two batches were held out for external validation purpose. The two subsets of INTERVAL data were put through the same quality control process as described below before performing training or validation. No significant technical variability was found between batches and hence batches within a subset (i.e. phase 1 or 2) were merged prior to the QC and genetic analysis including batch as a covariate to adjust for any residual batch effects. In the first step, samples with missing values for each of the ion-counts for a specific metabolite fragment (‘OrigScale’) were identified. These sample specific metabolite values were set to missing within the scaled and imputed data (‘ScaledImpData’), which contains for each metabolite the values within the OrigScale median normalised for run day (median set to 1 for run-day batch). Metabolites were then excluded if measured in only one batch or in less than 100 samples. Metabolite values were then winsorized to 5 standard deviation from the mean where the values exceeded mean +/-5 × standard deviation of the metabolite. Each metabolite was then log (natural) transformed prior to calculating the residuals adjusted for age, sex, Metabolon batch, INTERVAL recruitment centre, plate number, appointment month, the lag time between the blood donation appointment and sample processing, and the first 5 ancestry principal components. Prior to the genetic analysis, these residuals were standardised to a mean of 0 and standard deviation of 1. GWASs were then performed for each trait using the standardised trait values on samples of the first two batches, details of which were described in the previous study^47^. Finally, the standardised metabolites levels of the two INTERVAL subsets (batches 1+2 and batches 3+4) were further adjusted for the top 6-10 genetic principal components, which were used for genetic scores training and external validation respectively.

The Nightingale Health NMR platform was used to assay baseline serum samples of 45,928 INTERVAL participants and quantified 230 analytes in total, which are largely lipoprotein subfractions and ratios, lipids and low molecular weight metabolites. This study only focused on the 141 directly measured analytes and excluded those derived from other analytes. Apart from the missing values for low abundance analytes, the dataset also included zero values for some analytes, which were recoded as missing in our analysis. In addition, those analyte values of participants that had abnormally high/low values of more than 10 SD from the analyte mean across all participants were set as missing too. We further excluded participants with >30% analyte missingness and duplicate samples. Participants that failed genetic QC (see below) or did not have relevant phenotype data available were also removed, which resulted in 37,359 participants remaining in the analysis. Values of each analyte were log (natural) transformed and adjusted for age, sex, BMI, recruitment centre, time between blood draw and sample processing and the first 10 genetic principal component. The residuals were then inverse normal rank transformed, which were finally used to perform GWAS of these traits and their genetic score development. Details of quality control and GWAS for these traits can be found in the previous study^48^.

RNA sequencing was performed on the NovaSeq 6000 system (S4 flow cell, Xp workflow; Illumina) with 75 bp paired-end sequencing reads (reverse stranded) in INTERVAL, which were aligned to the GRCh38 human reference genome (Ensembl GTF annotation v99) using STAR (v2.7.3.a)^49^ and obtained the gene count matrix using featureCounts (v2.0.0)^50^. This in total resulted in raw gene-level count data of 60,676 genes (ENSEMBL gene IDs) across 4,778 individuals with 2.03–95.55 million uniquely mapped reads (median: ∼24 million). Poor-quality samples with RNA integrity number (RIN) < 4 or read depth < 10 million uniquely mapped reads were removed. We further removed one random individual from each flagged pair of related individuals, which were first-or second-degree estimated from genetic data. Finally, sample swaps and cross-contamination were assessed using match bam to VCF (MBV) method from QTLtools^51^, which identified and corrected 10 pairs of mislabelled samples; samples with no clear indication of their matching genotype data were also removed. Genes were retained based on >0.5 counts per million (CPM) expression threshold in ≥1% of the samples. The filtered count values were converted to trimmed mean of M-values (TMM)-normalized transcript per million mapped reads (FPKM) values^52^. Next, the normalised log_2_- FPKM values for each gene were ranked-based inverse normal transformed across samples. We further excluded globin genes, rRNA genes, and pseudogenes. After filtering, a total of 4,732 samples and 19,835 genes were retained for further eQTL analysis. Prior to eQTL mapping, the probabilistic estimation of expression residuals (PEER) method^53^ was used to find and correct for latent batch effects and other unknown confounders in the gene expression data. To estimate PEER factors independent of the effects of known variables, a set of 22 covariates of interest was included in the analysis. These were age, sex, BMI, and blood cell traits (N=19), including: (1) Basophil percentage (of white cell count); (2) Eosinophil percentage(of white cell count); (3) Lymphocyte percentage (of white blood cell count; (4) Monocyte percentage (of white blood cell count); (5) Neutrophil percentage (of white blood cell count); (6) White blood cell (leukocyte) count (reported); (7) Immature reticulocyte fraction; (8) Haematocrit (volume percentage of blood occupied by red cells); (9) Reticulocyte percentage (of red cell and reticulocyte count); (10) Haemoglobin concentration; (11) Mean corpuscular haemoglobin; (12) Mean corpuscular haemoglobin concentration; (13) Mean corpuscular (red cell) volume; (14) Red blood cell (erythrocyte) count (reported); (15) Red cell distribution width; (16) Mean platelet volume; (17) Plateletcrit; (18) Platelet distribution width; (19) Platelet count. The eQTL mapping was performed on genome-wide variants using TensorQTL v1.0.3^54^ adjusting for age, sex, BMI, the above mentioned blood cells traits (N=19), the top 10 genetic principal components, RIN, sequencing batch, RNA concentration, raw read depth, season (based on month of blood draw), and PEER factors (N=30). The normalised gene level values were also adjusted for the same set of covariates used in the eQTL mapping for their genetic score training and validation. Note that we held out the last two batches of samples for external validation purpose and the first four were used for eQTL mapping and genetic score training/internal validation.

The genotyping and its quality control for INTERVAL samples have been previously described in detail^55^. The samples were genotyped using the Affymetrix UK Biobank Axiom array, which assays approximately 830,000 variants. The variants were phased using SHAPEIT3 and imputed on a combined 1000 Genomes Phase 3-UK10K reference panel. After various quality control steps, it finally results in 10,572,788 variants for 43,059 samples. The number of valid samples in each platform for genetic score construction (**Table 1)** excluded samples that did not pass the genetic QC.

### External validation cohorts

The FENLAND study profiled the plasma proteins of 12,084 participants using the aptamer-based SomaScan assay (version 4), in which 8994 participants were genotyped using the same the Affymetrix UK Biobank Axiom array as INTERVAL^43^. The later subset of Fenland participants were used for the genetic score model validation in our study. As FENLAND and INTERVAL applied two different versions of the SomaScan array (versions 3 and 4), we matched aptamers (or SOMAmers) between the two studies by using their unique SomaScan IDs, which resulted in 2129 matched results. The detailed QC steps for protein measurements, and genotype imputation and QC for genotype data in the FENLAND study were described in the previous study^19^. The Fenland study was approved by the National Health Service (NHS) Health Research Authority Research Ethics Committee (NRES Committee – East of England Cambridge Central, ref. 04/Q0108/19), and all participants provided written informed consent. Both the Orkney Complex Disease Study (ORCADES)^22^ and Northern Sweden Population Health Study (NSPHS)^20^ have measured plasma protein levels of their participants on the four Olink panels that were used in INTERVAL, and genotyped participants using Illumina arrays. Thus, participants of the two studies were used to validate genetic score models of Olink proteins considered in our study, where gene names of proteins were used to match proteins between studies. For those proteins that appeared in two or more Olink panels, their validation measurements were averaged across panels for the protein. Detailed imputation and QC steps for protein abundance measurements and genetic data in the two studies were described in the previous studies^56,57^. Protein levels in ORCADES were adjusted for age, sex, plate, plate row, and plate column, sampling year and season, top 10 genetic PCs and kinship before used for validation. ORCADES also used the same platform Metabolon HD4 as INTERVAL to measure plasma metabolites of participants, and we used COMP identifier in the platform to match metabolites between the two studies, which resulted in 455 overlapped traits. Detailed quality control steps for metabolites in ORCADES were described in the previous study^23^ and their trait levels were adjusted for covariates of sex, age, BMI, sampling season and year, plate number, plate column, plate row, genotyping array and top 20 PCs. The UK Biobank, ORCADES and the VIKING health study^26^ were used as external cohorts to validate genetic scores of Nightingale traits, and traits identifiers provided in the platform were used to successfully match all 141 traits between these studies and INTERVAL. Quality control for these traits in each external cohort has been described previously in details^23,24^. Before validation, levels of these traits were adjusted for sex, age, BMI, sampling season and sampling year, genotyping array and top 20 genetic PCs in ORCADES, VIKING; in UKB, they were adjusted for sex age, BMI, use of lipid lowering medication, top 10 genetic PCs and technical variance following the protocol of the previous study^24^.

The Multi-Ethnic Cohort (MEC) recruited three major Asian ethnic groups represented in Singapore: Chinese, Malays and Indians, between 2004 and 2010 to better understand how genes and lifestyle influence health and diseases differently in persons of different ethnicities^28^. Between 2011 and 2016, the participants were further invited for a follow-up. Whole genome-sequencing was performed on 2,902 MEC participants as Phase I of the National Precision Medicine Programme (https://npm.a-star.edu.sg/). Samples were whole-genome sequenced to an average of 15X coverage. Read alignment was performed with BWA-MEM and variant discovery and genotyping was performed with GATK. Site-level filtering includes only retaining VQSR-PASS and non-STAR allele variants. At the sample level, samples with call rate <95%, BAM cross-contamination rate >2%, or BAM error-rate > 1.5%; at the genotype call level, genotypes with DP<5 or GQ<20 or AB>0.8 (heterozygotes calls), were set to NULL. Finally, samples with abnormal ploidy were excluded, and genetic ancestry were determined with k-means clustering from the top 15 principal components. Both SomaScan (version 4) and Nightingale NMR platforms were used to assay baseline and revisit blood samples of participants in MEC. For quality control of Nightingale data, participants with >10% missing metabolic biomarker values were excluded from subsequent analyses. For participants with biomarker values lower than detection level, we replaced values of 0 with a value equivalent to 0.9 multiplied by the non-zero minimum value of that measurement. For quality control of SomaScan data, protein levels were first normalized to remove hybridization variation within a run. This was followed by median normalization across calibrator control samples to remove other assay biases within the run. Overall scaling and calibration were then performed on a per-plate basis to remove overall intensity differences between runs with calibrator controls. Finally, median normalization to a reference was performed on the individual samples with QC controls. During these standardization steps, multiple scaling factors were generated for each sample/aptamer at each step. The final number of samples in each ethnic groups used in our validation were given in **Table 1**. For both SomaScan and Nightingale traits, natural log-transformation was applied before adjusting for age, sex, T2D status, and BMI (Nightingale traits only). Residuals from the regression were inverse-normalised for correlation analyses with genetic scores trained in INTERVAL.

The Jackson Heart Study (JHS) is a community-based longitudinal cohort study begun in 2000 of 5,306 self-identified Black individuals from the Jackson, Mississippi metropolitan statistical area^29,58^. The participants included in our validation of genetic scores for SomaScan proteins are samples collected at Visit 1 between 2000 and 2004 from 1,852 individuals with whole genome sequencing and proteomic profiling (SomaScan) performed, quality controls of which were detailed in the previous studies^29,59,60^. SomaScan IDs were used to match shared proteins between JHS and INTERVAL, which identified 820 proteins in total. Protein levels were adjusted for age, sex and the first 10 principal components of genetic ancestry in JHS, before they were used for evaluating performance of genetic scores.

### Polygenic scoring method

A genetic score is most commonly constructed as a weighted sum of genetic variants carried by an individual, where the genetic variants are selected and their weights quantified via univariate analysis in a corresponding genome-wide association study^61,62^:

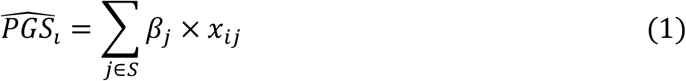

where *S* is the set of variants, referring to single nucleotide polymorphisms (SNPs) in this study, that are identified in the variant selection step described below; *β*^*j*^ *is* the effect size of the SNP *j* that is obtained through the univariate statistical association tests in the GWAS; *x*^*ij*^ is the genotype dosage of SNP *j* of the individual *i*. As the variant set *S* is derived through a LD pruning and p-value thresholding process, this method is often named as the P+T. However, P+T relies on hard cut-off thresholds to remove LD correlations among variants and select associated variants. It is often challenging to balance between keeping predictive variants and removing redundant and uninformative variants that can limit the prediction precision. Also, due to the inherent linear assumption of the univariate analysis in P+T, this method leaves no modelling considerations for joint effects between variants. To alleviate these limitations, various machine learning based methods, such as Bayesian ridge (BR), elastic net (EN)^45^ and LDpred^63^, have been utilized to construct genetic scores for a wide range of traits and diseases^8^. In particular, BR and EN have been shown to outperform other methods when developing scores for predicting biomolecular traits, such as blood cell traits and gene expression^8,10^, which are similar to the type of traits considered in this study. We adopted the BR method for the genetic score construction of all the biomolecular traits as BR is more efficient to run in practice (see details below).

Bayesian ridge is a multivariate linear model which assumes that the genetic variants have linear additive effects on the genetic score of the trait^8,64^. In addition, BR also assumes that the genetic score of a trait follows a Gaussian distribution, and the prior for effect sizes of variants is also given by a spherical Gaussian:

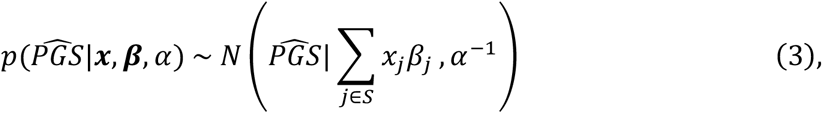

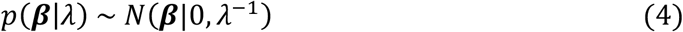

where α and λ are coefficients of the model and subject to two Gamma distribution: Gamma(α_1_, α_2_) and Gamma(λ_1_, λ_2_). These two prior Gamma distributions can be set via a validation step.

### Genetic score model training and evaluation

The explained variance (R^*2*^) and Spearman’s rank correlation coefficient were used to measure the performance of constructed genetic scores in the INTERVAL training samples and external cohorts (or INTERVAL withheld subset), where R^*2*^ scores were calculated using the squared Pearson correlation coefficient. We adopted a similar strategy for sample partition when training and evaluating genetic scores within the training samples as previous studies^8,10^ that utilised learning-based methods to construct genetic scores for molecular traits. The training samples of a trait were randomly and equally partitioned to five subsets, from which any four subsets are used as true-training data to learn a genetic score model of the trait, and test the model’s performance on the remaining 20% of samples. Given a genetic scoring method and a trait, we obtained five different genetic score models of the trait and the mean of their performance measurements in the corresponding testing samples in INTERVAL was reported (internal validation). Note that, due to the high similarities between the five genetic score models trained for most traits, only one model was randomly selected from the five and evaluated in the external cohorts (or INTERVAL withheld subset).

When training genetic score models using the BR method, we need to select two appropriate prior gamma distributions, i.e. α_1_, α_2_, λ_1_ and λ_2_. To do so, a grid search across the set [-10^10^, - 10^5^,-10, 0, 10, 10^5^, 10^10^] was performed on the true-training data set, in which 10% of the samples were used as a validation set. However, running this validation process is resource and time-intensive, which makes it challenging to run for all the traits. To address this problem, we found that it is reasonable to assume that the same category of molecular traits, i.e. proteomic traits or metabolomic traits, share the same prior distributions, without sacrificing model performance. Thus, we only needed to run the validation process once for each of the platforms (a trait was randomly selected), and applied the identified optimal prior distributions to other traits.

### Variant selection and performance comparison between BR and P+T

Selecting a proper set of variants and feeding into a polygenic scoring method are a key step for effective genetic score construction. To do so and further confirm the superiority of BR method, we looked at the performance of BR and P+T on a variety of variant selection schemes for the traits in three platforms (SomaScan, Olink and Metabolon).

To ensure the generalizability of genetic score models when applied to other cohorts, a variant filtering step was first performed for all the traits considered, which applied a MAF threshold of 0.5% and excluded all multi-allelic variants as well as ambiguous variants (i.e. A/T, G/C). To remove LD dependencies among variants, a follow-up LD thinning step was carried out at an *r*^*2*^ threshold of 0.8 on all the variants. The remaining variants were then filtered at given p-value thresholds (from their GWAS summary statistics conducted on the INTERVAL training data) for a trait in different platforms. To identify an appropriate variant selection scheme for the use of all the biomolecular traits, we attempted the following four p-value thresholding schemes for protein traits in Olink and SomaScan platforms: (1) p-value < 5 × 10^−8^ on all the variants; (2) p-value < 5 × 10^−8^ on variants in the *cis* region only (within 1MB of the corresponding gene’s transcription start site); (3) all the *cis* variants only; (4) all the *cis* variants and p-value < 1 × 10^−3^ on the *trans* variants; and the two different p-value thresholds on the genome-wide variants for metabolite traits in the Metabolon platform (as they do not distinguish *cis* and *trans* regions): (1) p-value < 5 × 10^−8^; (2) p-value < 1 × 10^−3^.

Then, we compared the performance of BR and P+T on these variant sets in the internal validation (**Figure S1-S3**). Regarding the proteomic traits (SomaScan and Olink), the two variant selection schemes: (1) p-value < 5×10^−8^ on genome-wide variants and (2) all the *cis* variants and p-value < 1×10^−3^ on the *trans* variants, were shown to be the best performing schemes with either of the methods; BR method largely outperformed P+T across the two variant selection schemes. Meanwhile, it was noted that the two selection schemes led to greatly different performance, with the latter scheme achieving an unrealistic mean R^*2*^ of ∼0.74 across all the proteins (only ∼0.09 for the former scheme). Similarly, for the metabolomic traits (Metabolon), the applied two variant selection schemes significantly differ in their performance in internal validation, and BR was also shown to a better performing method.

To further identify the optimal variant selection scheme, we also looked at the performance of validated genetic score models trained with the two identified (for proteins) or all the two applied (for metabolites) schemes using BR method for Olink traits and Metabolon traits (**Figure 3** and **Figure S4**) in external cohorts (NSPHS and ORCADES) or withheld INTERVAL data. Despite the second scheme (all the *cis* variants and p-value < 1×10^−3^ on the *trans* variants for proteins, or p-value < 1×10^−3^ on genome-wide variants for metabolites) showed outstanding performance in internal validation, its performance saw a dramatic decline in external validation for almost every trait validated (**Figure S4**). It indicates this variant selection scheme caused an overfitting problem in genetic score training, which is consistent with previous findings when using overly lenient p-value thresholds for variant selection^8^. These results suggested that the BR method with the variant selection scheme of p-value < 5×10^−8^ on genome-wide variants was the optional method (of those tested) for genetic score development of these biomolecular traits, thus we applied this approach to all other traits for their genetic score development in this study.

### Comparing the genetic scores for shared proteins between SomaScan and Olink

SomaScan and Olink used two different technologies for protein level measurement. The two platforms measured many proteins in common,among which there are 169 unique proteins whose genetic scores we have validated. To check the impact of technologies on genetic prediction, we looked at how the genetic scores trained on one platform can predict protein levels from the other platform on the INTERVAL training samples (**Figure S16)**. We confirmed that performance of these overlapped genetic scores trained in the other platform was generally consistent with that of the scores trained in their original platform. However, we did observe, in some cases, the genetic scores trained in the two platforms can lead to very different predictions, for which we found that they are mainly due to the differences in what the two platforms are actually quantifying. For example, among the 169 proteins, there are 11 proteins in SomaScan that had a R^*2*^ > 0.3 in internal validation, in which 10 proteins also achieved a R^*2*^ > 0.3 but the remaining protein (CHI3L1) received a poor R^*2*^ < 0.1 when predicting with Olink genetic scores. We found that the remaining protein received a lowest Pearson’s *r* score among the 11 proteins between their actual protein levels measured in the two platforms. In INTERVAL, there were ∼700 participants (depending on the protein) who were assayed by both SomaScan and Olink, which allowed us to calculate the correlations between the actual protein levels measured by the two platforms for the same protein. These results suggested, despite great consistency, genetic scores of the same protein trained in the two platforms can represent distinct aspects of protein biology of prediction and integration of diverse proteomic techniques may enable to develop better genetic scores for these proteins^65^.

### Pathway coverage analysis of heritable proteins

In this analysis, SomaScan and Olink proteins were combined based on their Uniprot ID, where duplicate proteins were removed if identified. We only kept proteins with R^*2*^ > 0.01 in internal validation, resulting in a total of 2,205 unique proteins for the analysis. We used pathway data of Homo sapiens curated at Reactome^30^ and conducted analyses to uncover the coverage of these proteins in the pathways. In detail, this analysis looked at the percentages of these proteins in annotated physical entities of each super-pathway, and the percentages of the lowest-level pathways these proteins covered among all the lowest-level pathways of each super-pathway. Where at least one protein in this study are included in entities of a lowest-level pathway, we considered this pathway is covered by proteins of this study.

### Phenome-wide association analysis (PheWAS) in UKB

We included biomolecular traits with *R*^*2*^ > 0.01 in internal validation in this analysis (11,576 traits in total) and considered only participants of European ancestry in UKB (the White British subset). We used the version 3 of imputed and quality controlled genotype data for UKB, which were detailed in the previous study^25^. Using version 1.2 of the PheWAS Catalog^31^, we extracted the curated phenotype definitions of all phecodes. Each phecode is provided as a set of WHO International Classification of Diseases (ICD) diagnosis codes in versions 9 (ICD-9) and 10 (ICD-10) of the ontology to define individuals with the phenotype of interest, and a set of related phecodes that should be excluded from the control cohort of unaffected individuals. To define cases for each phecode, we searched for the presence of any of the constituent ICD-9/10 codes in linked health records (including in-patient Hospital Episode Statistics data, cases of invasive cancer defined in the cancer registry, and primary and secondary cause of death information from the death registry), and converted the earliest coded date to the age of phenotype onset. Individuals without any codes for the phenotype of interest were recorded as controls, and censored according to the maximum follow-up of the health linkage data (January 31, 2020) or the date of death whichever came first. To define the cohort for testing molecular genetic score associations with the age-of-onset of each phenotype, we used the set of events and censored individuals described above and removed any individuals with related phenotypes (based on definitions from the PheWAS Catalog), restricting analyses to be sex-specific (e.g. ovarian and prostate cancer) where requires. To ensure a well-powered study we restricted the PheWAS analysis to phenotypes with at least 200 cases in the 409,703 European ancestry individuals whose reported sex match the genetically inferred sex from the UKB quality controlled genotype data^25^, resulting in a set of 1,123 phecodes included in the final analysis. The association of the genetic score for biomolecular traits with the onset of each phenotype was assessed by using a Cox proportional hazards model with age-as-timescale, stratified by sex and adjusted for genotyping array and 10 PCs of genetic ancestry. The association between genetic scores and each phecode is reported in terms of its effect size (Hazard ratio) and corresponding significance (p-value), and significant results were defined as Benjamini/Hochberg FDR-corrected p-value < 0.05 for all the tested traits. Statistical analyses were performed in python and the Cox model was implemented using the lifelines package^66^.

## Acknowledgement

Participants in the INTERVAL randomised controlled trial were recruited with the active collaboration of NHS Blood and Transplant England (www.nhsbt.nhs.uk), which has supported field work and other elements of the trial. DNA extraction and genotyping were co-funded by the National Institute for Health Research (NIHR), the NIHR BioResource (http://bioresource.nihr.ac.uk) and the NIHR Cambridge Biomedical Research Centre (BRC-1215-20014) [*]. The academic coordinating centre for INTERVAL was supported by core funding from the: NIHR Blood and Transplant Research Unit in Donor Health and Genomics (NIHR BTRU-2014-10024), UK Medical Research Council (MR/L003120/1), British Heart Foundation (SP/09/002; RG/13/13/30194; RG/18/13/33946) and NIHR Cambridge BRC (BRC-1215-20014) [*]. A complete list of the investigators and contributors to the INTERVAL trial is provided in reference [**]. The academic coordinating centre would like to thank blood donor centre staff and blood donors for participating in the INTERVAL trial. UK Biobank data access was approved under project 7439, and all the participants gave their informed consent for health research. Generation of part of the Metabolon data in INTERVAL was funded by Biomarin Pharmaceuticals. The Multi-Ethnic Cohort (MEC) is funded by individual research and clinical scientist award schemes from the Singapore National Medical Research Council (NMRC, including MOH-000271-00) and the Singapore Biomedical Research Council (BMRC), the Singapore Ministry of Health (MOH), the National University of Singapore (NUS) and the Singapore National University Health System (NUHS). This work on omics polygenic score transferability is supported by the NUS-Cambridge Seed Grant July 20201 (NUSMEDIR/Cambridge/2021-07/001). The metabolite biomarkers data were generated in collaboration with Nightingale Health Ltd. The protein biomarker data were generated in collaboration with Somalogic Inc. The MEC whole genome sequence data made use of data generated as part of the Singapore National Precision Medicine (NPM) program funded by the Industry Alignment Fund (Pre-Positioning) (IAF-PP: H17/01/a0/007). NPM made use of data/samples collected in the following cohorts in Singapore: (1) The Health for Life in Singapore (HELIOS) study at the Lee Kong Chian School of Medicine, Nanyang Technological University, Singapore (supported by grants from a Strategic Initiative at Lee Kong Chian School of Medicine, the Singapore Ministry of Health (MOH) under its Singapore Translational Research Investigator Award (NMRC/STaR/0028/2017) and the IAF-PP: H18/01/a0/016); (2) The Growing up in Singapore Towards Healthy Outcomes (GUSTO) study, which is jointly hosted by the National University Hospital (NUH), KK Women’s and Children’s Hospital (KKH), the National University of Singapore (NUS) and the Singapore Institute for Clinical Sciences (SICS), Agency for Science Technology and Research (A*STAR) (supported by the Singapore National Research Foundation under its Translational and Clinical Research (TCR) Flagship Programme and administered by the Singapore Ministry of Health’s National Medical Research Council (NMRC), Singapore-NMRC/TCR/004-NUS/2008; NMRC/TCR/012-NUHS/2014. Additional funding is provided by SICS and IAF-PP H17/01/a0/005); (3) The Singapore Epidemiology of Eye Diseases (SEED) cohort at Singapore Eye Research Institute (SERI) (supported by NMRC/CIRG/1417/2015; NMRC/CIRG/1488/2018; NMRC/OFLCG/004/2018); (4) The Multi-Ethnic Cohort (MEC) cohort (supported by NMRC grant 0838/2004; BMRC grant 03/1/27/18/216; 05/1/21/19/425; 11/1/21/19/678, Ministry of Health, Singapore, National University of Singapore and National University Health System, Singapore); (5) The SingHealth Duke-NUS Institute of Precision Medicine (PRISM) cohort (supported by NMRC/CG/M006/2017_NHCS; NMRC/STaR/0011/2012, NMRC/STaR/ 0026/2015, Lee Foundation and Tanoto Foundation); (6) The TTSH Personalised Medicine Normal Controls (TTSH) cohort funded (supported by NMRC/CG12AUG17 and CGAug16M012). The views expressed are those of the author(s) are not necessarily those of the National Precision Medicine investigators, or institutional partners. We are grateful to all Fenland volunteers and to the General Practitioners and practice staff for assistance with recruitment. We thank the Fenland Study Investigators, Fenland Study Co-ordination team and the Epidemiology Field, Data and Laboratory teams. Proteomic measurements were supported and governed by a collaboration agreement between the University of Cambridge and SomaLogic. The Fenland Study (10.22025/2017.10.101.00001) is funded by the Medical Research Council (MC_UU_12015/1). We further acknowledge support for genomics from the Medical Research Council (MC_PC_13046). The Orkney Complex Disease Study (ORCADES) was supported by the Chief Scientist Office of the Scottish Government (CZB/4/276, CZB/4/710), a Royal Society URF to J.F.W., the MRC Human Genetics Unit quinquennial programme “QTL in Health and Disease”, Arthritis Research UK and the European Union framework program 6 EUROSPAN project (contract no. LSHG-CT-2006-018947). DNA extractions were performed at the Edinburgh Clinical Research Facility, University of Edinburgh. We would like to acknowledge the invaluable contributions of the research nurses in Orkney, the administrative team in Edinburgh and the people of Orkney. The Viking Health Study – Shetland (VIKING) was supported by the MRC Human Genetics Unit quinquennial programme grant “QTL in Health and Disease”. DNA extractions and genotyping were performed at the Edinburgh Clinical Research Facility, University of Edinburgh. We would like to acknowledge the invaluable contributions of the research nurses in Shetland, the administrative team in Edinburgh and the people of Shetland. We acknowledge support from the MRC Human Genetics Unit programme grant, “Quantitative traits in health and disease” (U. MC_UU_00007/10). Whole genome sequencing (WGS) for the Trans-Omics in Precision Medicine (TOPMed) program was supported by the National Heart, Lung and Blood Institute (NHLBI). WGS for “NHLBI TOPMed: Jackson Heart Study” (phs000964) was performed at the Northwest Genomics Center (HHSN268201100037C). Core support including centralized genomic read mapping and genotype calling, along with variant quality metrics and filtering were provided by the TOPMed Informatics Research Center (3R01HL-117626-02S1; contract HHSN268201800002I). Core support including phenotype harmonization, data management, sample-identity QC, and general program coordination were provided by the TOPMed Data Coordinating Center (R01HL-120393; U01HL-120393; contract HHSN268201800001I). We gratefully acknowledge the studies and participants who provided biological samples and data for TOPMed. The Jackson Heart Study (JHS) is supported and conducted in collaboration with Jackson State University (HHSN268201800013I), Tougaloo College (HHSN268201800014I), the Mississippi State Department of Health (HHSN268201800015I) and the University of Mississippi Medical Center (HHSN268201800010I, HHSN268201800011I and HHSN268201800012I) contracts from the National Heart, Lung, and Blood Institute (NHLBI) and the National Institute on Minority Health and Health Disparities (NIMHD). The authors also wish to thank the staffs and participants of the JHS. JHS disclaimer - The views expressed in this manuscript are those of the authors and do not necessarily represent the views of the National Heart, Lung, and Blood Institute; the National Institutes of Health; or the U.S. Department of Health and Human Services. YX and MI were supported by the UK Economic and Social Research Council (ES/T013192/1). SCR is funded by a BHF Programme Grant (RG/18/13/33946). CL, MP, JL are funded by the Medical Research Council (MC_UU_00006/1 - Aetiology and Mechanisms). JD holds a British Heart Foundation Professorship and a NIHR Senior Investigator Award [*]. MI is supported by the Munz Chair of Cardiovascular Prediction and Prevention and the NIHR Cambridge Biomedical Research Centre (BRC-1215-20014) [*]. This study was supported by the Victorian Government’s Operational Infrastructure Support (OIS) program. We acknowledge Ben Sun and Tao Jiang for previous analyses of INTERVAL SomaScan and genotype quality control, respectively. For the purpose of open access, the author has applied a Creative Commons Attribution (CC BY) licence to any Author Accepted Manuscript version arising from this submission.

*The views expressed are those of the author(s) and not necessarily those of the NHS, the NIHR or the Department of Health and Social Care. **Di Angelantonio E, Thompson SG, Kaptoge SK, Moore C, Walker M, Armitage J, Ouwehand WH, Roberts DJ, Danesh J, INTERVAL Trial Group. Efficiency and safety of varying the frequency of whole blood donation (INTERVAL): a randomised trial of 45,000 donors. Lancet. 2017 Nov 25;390(10110):2360-2371.

