## Supplementary Figures for "An atlas of genetic scores to predict multi-omic traits"

**Figure S1: Performance ($R^2$) distribution of genetic scores and performance comparison ($R^2$) between Bayesian Ridge (BR) and pruning and thresholding (P+T) methods for Metabolon traits in internal validation.** The density plots show the distributions of $R^2$ performance for genetic scores developed using BR method on different variant sets. P+T constructs genetic scores using weighted sum of a selected genetic variant set, where GWAS effect sizes of these variants are used as their weights. The scatter plots compare the performance of genetic scores developed using BR and P+T on different variant sets. It is noted that the variant set with p-value <1e-3 resulted in an overfitting problem (see **Figure S4** for details).

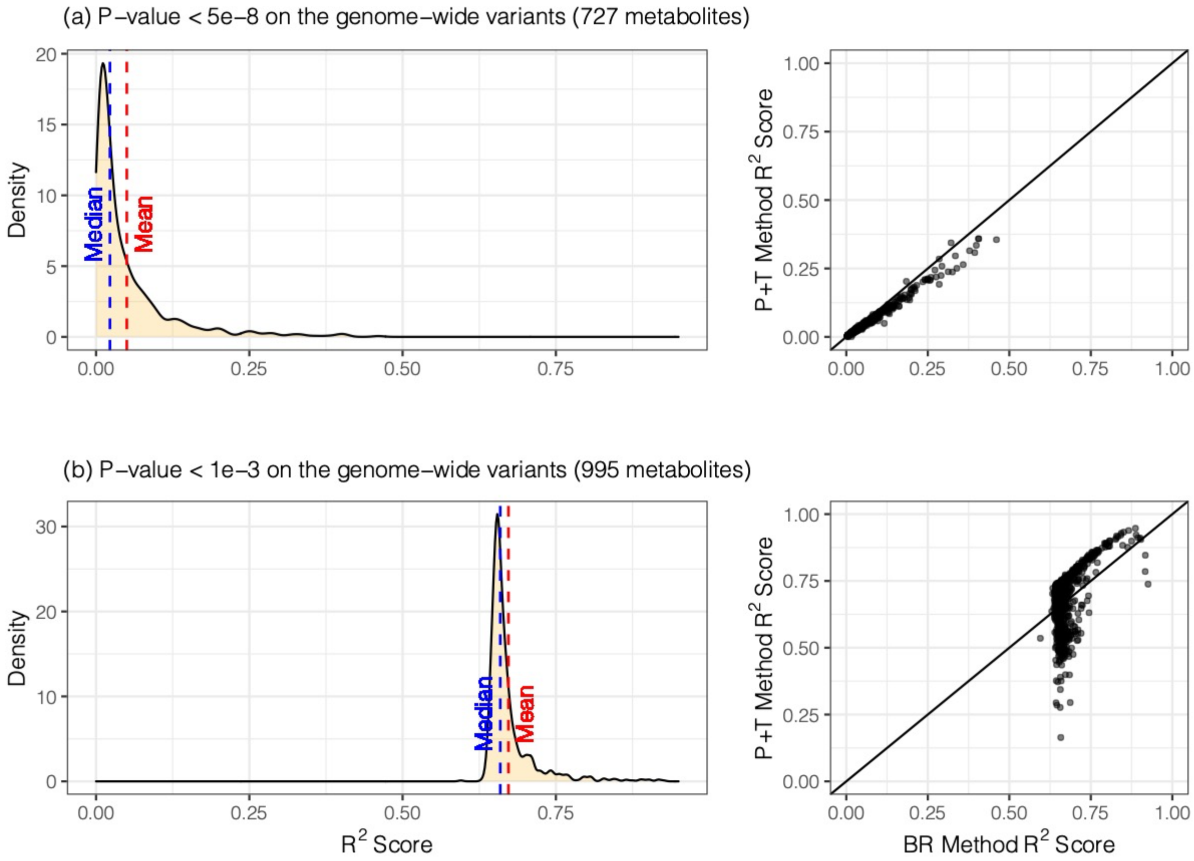

**Figure S2: Performance ($R^2$) distribution of genetic scores and performance comparison ($R^2$) between BR and P+T methods for Olink traits in internal validation.** The density plots show the distributions of $R^2$ performance for genetic scores developed using BR method on different variant sets. The scatter plots compare the performance of genetic scores developed using BR and P+T on different variant sets.

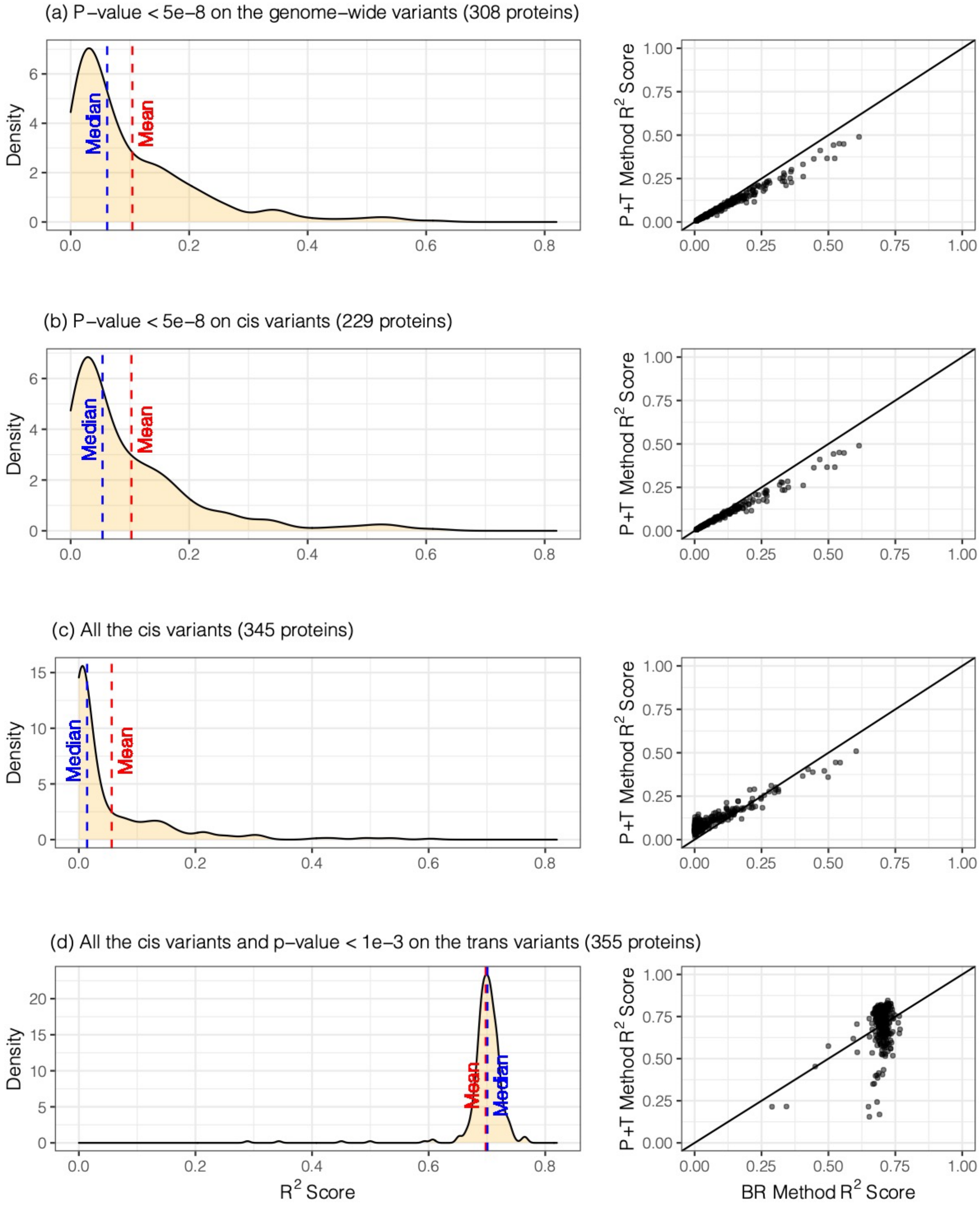

**Figure S3: Performance ($R^2$) distribution of genetic scores and performance comparison ($R^2$) between BR and P+T methods for SomaScan traits in internal validation.** The density plots show the distributions of $R^2$ performance for genetic scores developed using BR method on different variant sets. The scatter plots compare the performance of genetic scores developed using BR and P+T on different variant sets.

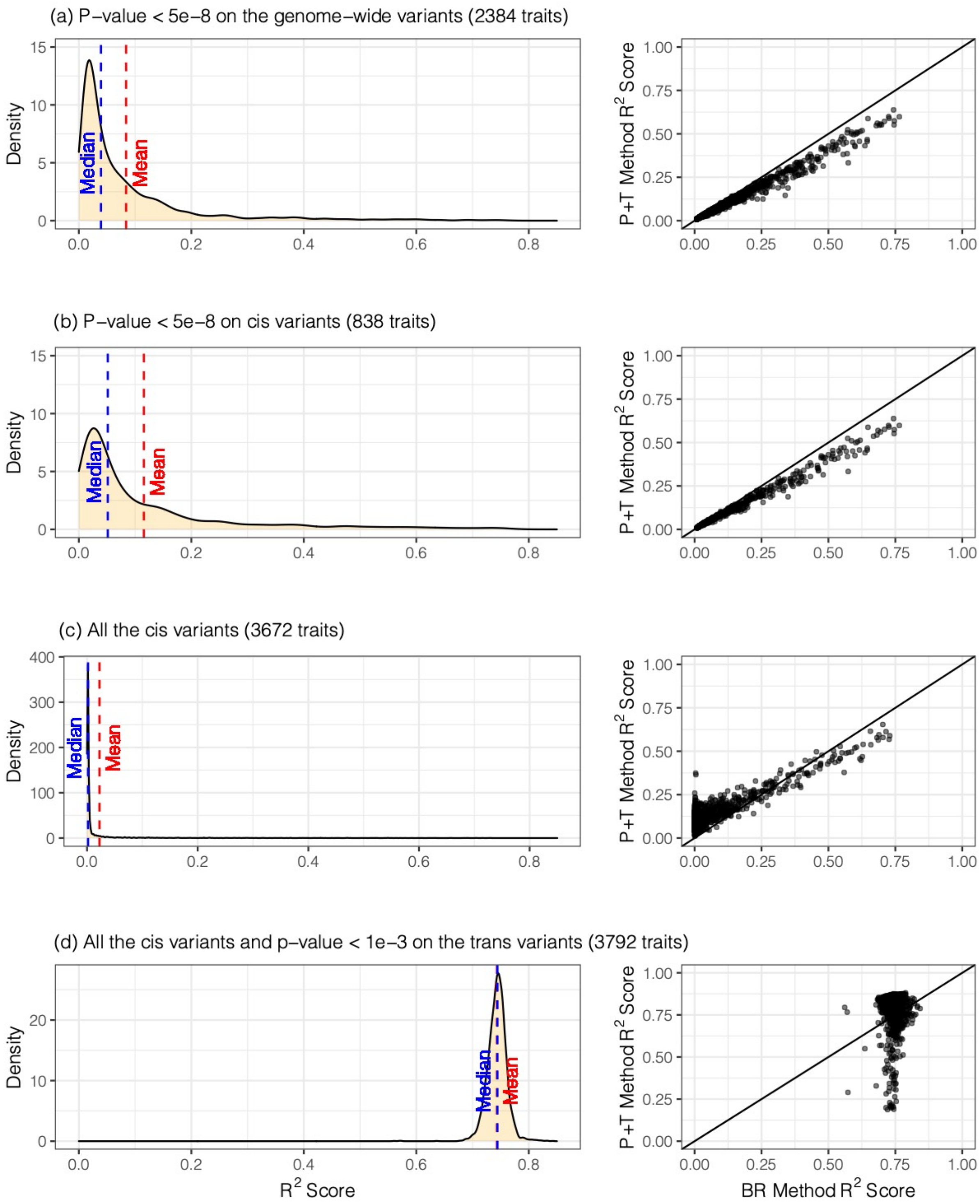

**Figure S4: Performance ($R^2$) comparison between internal and external validation for genetic scores of Metabolon and Olink traits.** The genetic scores were constructed using BR method on the set of genome wide variants with p-value< $1\times10^{-3}$.

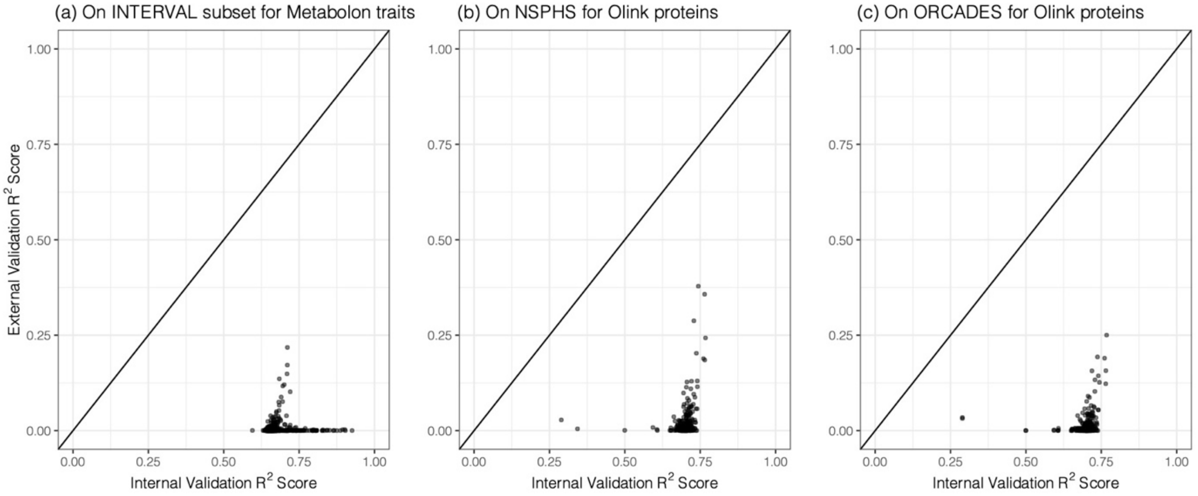

**Figure S5: Distribution of the number of variants in the genetic scores and the correlations between performance ($R^2$) of genetic scores and the number of variants comprising the score.** The density plots show the distribution of number of variants comprising the genetic scores at each platform. The scatter plots show the change of $R^2$ score in the internal validation by the number of variants in the genetic score model.

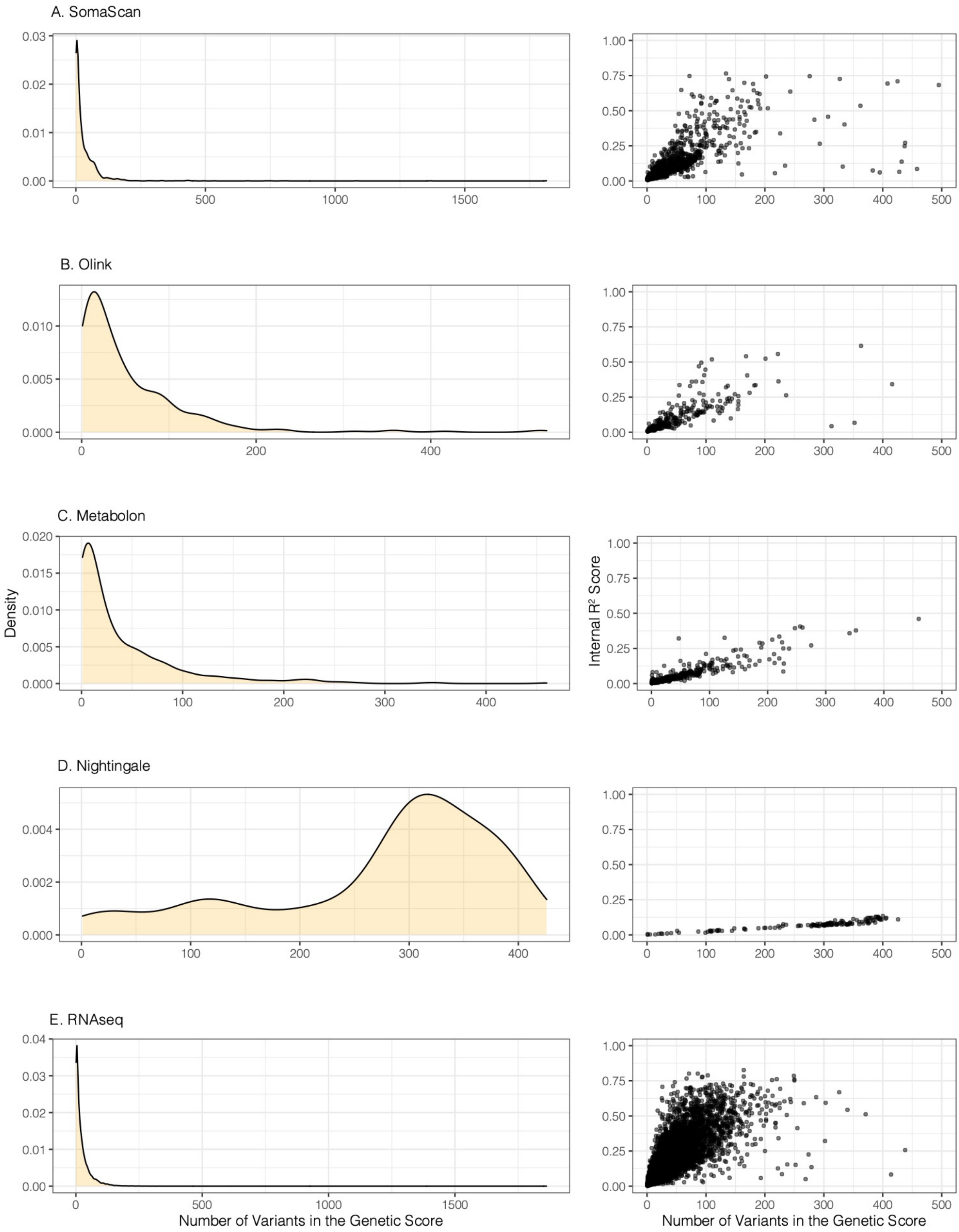

**Figure S6: Distribution of $R^2$ performance in external validation for genetic scores of Metabolon traits.** This analysis included all the traits validated in the external cohort or the INTERVAL withheld set.

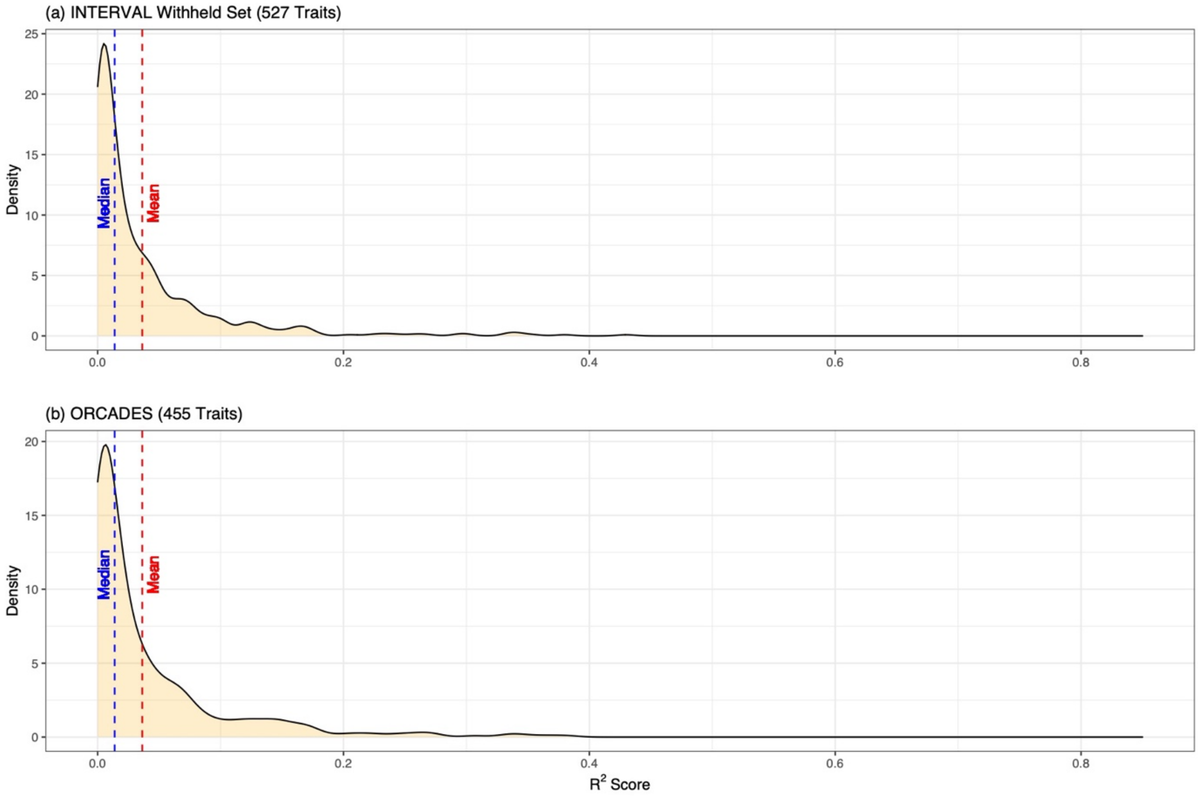

**Figure S7: Distribution of $R^2$ performance in external validation for genetic scores of Nightingale traits.** This analysis included all the traits validated in each external cohort.

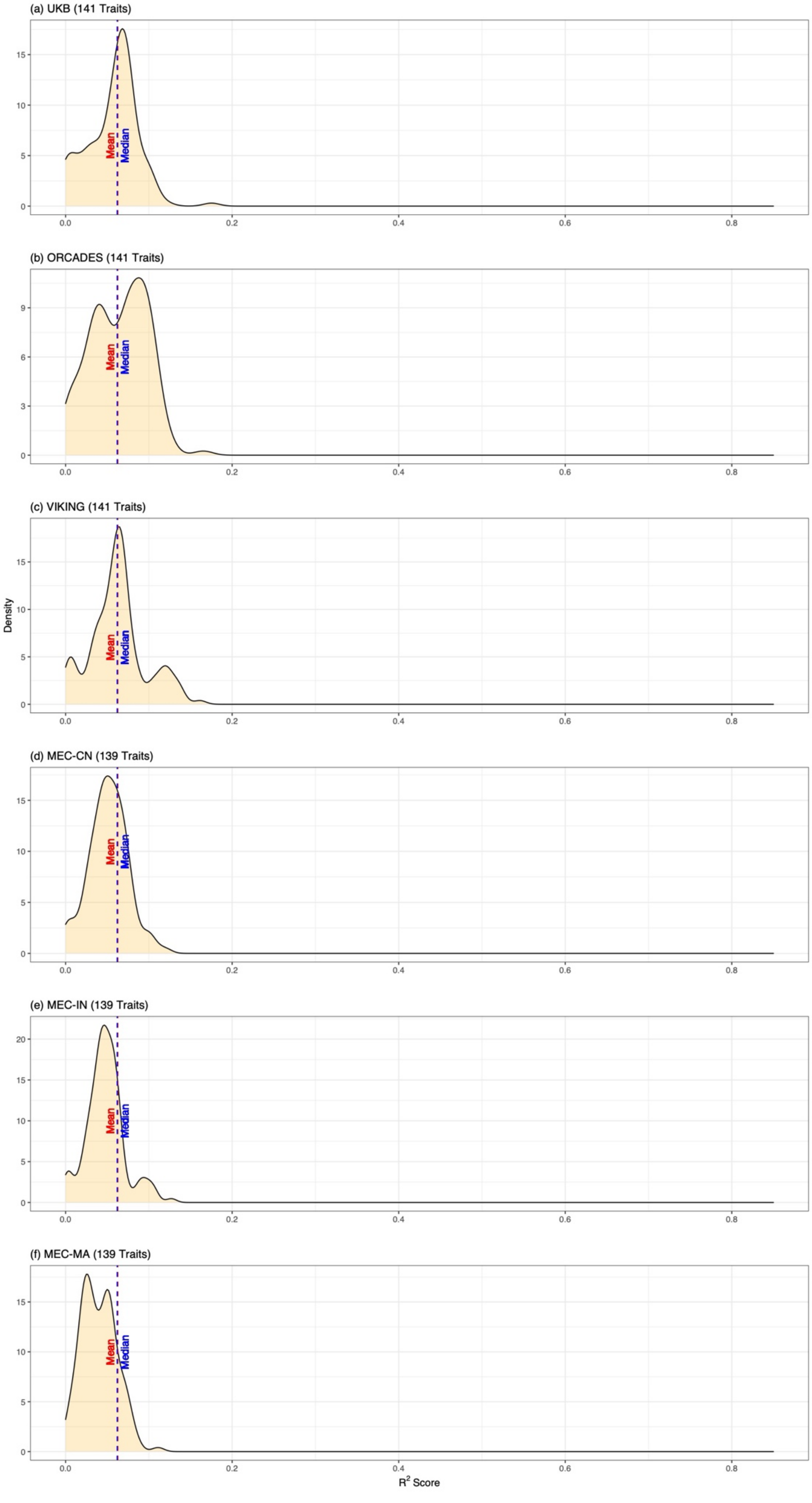

**Figure S8: Distribution of $R^2$ performance in external validation for genetic scores of Olink traits.** This analysis included all the traits validated in each external cohort.

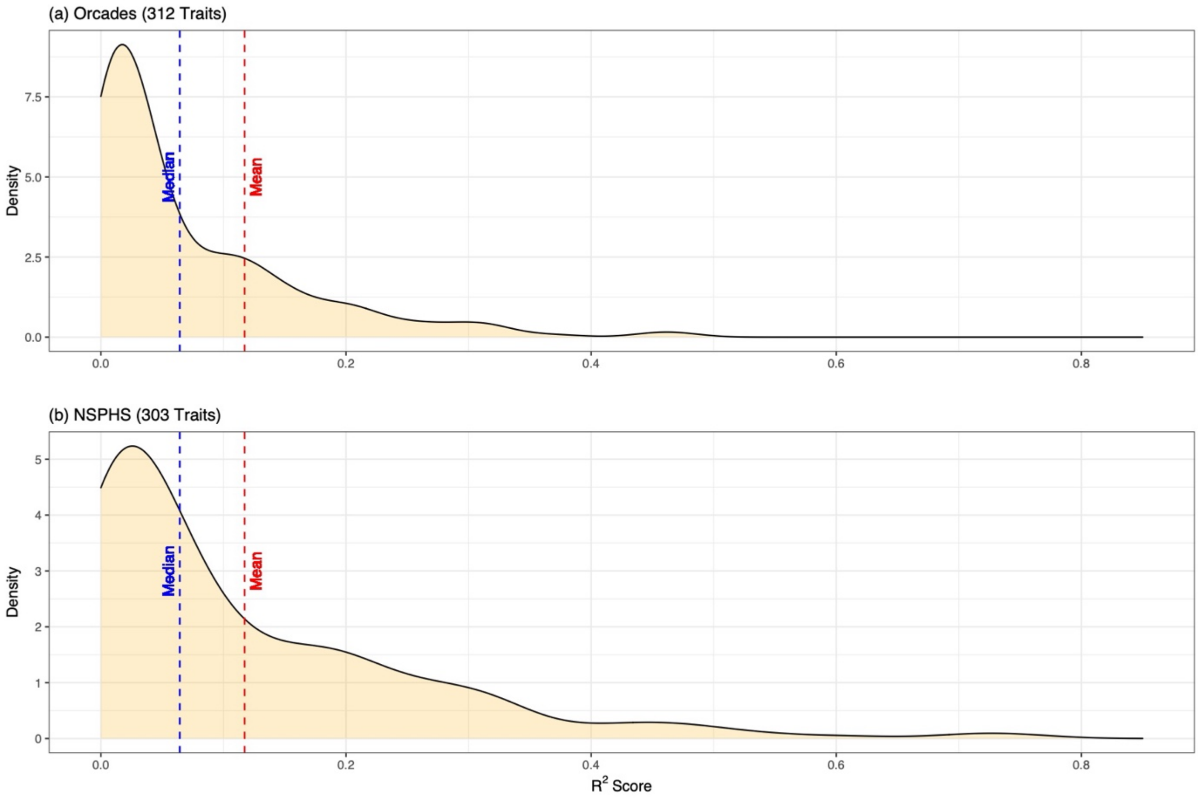

**Figure S9: Distribution of $R^2$ performance in external validation for genetic scores of SomaScan traits.** This analysis included all the traits validated in each external cohort.

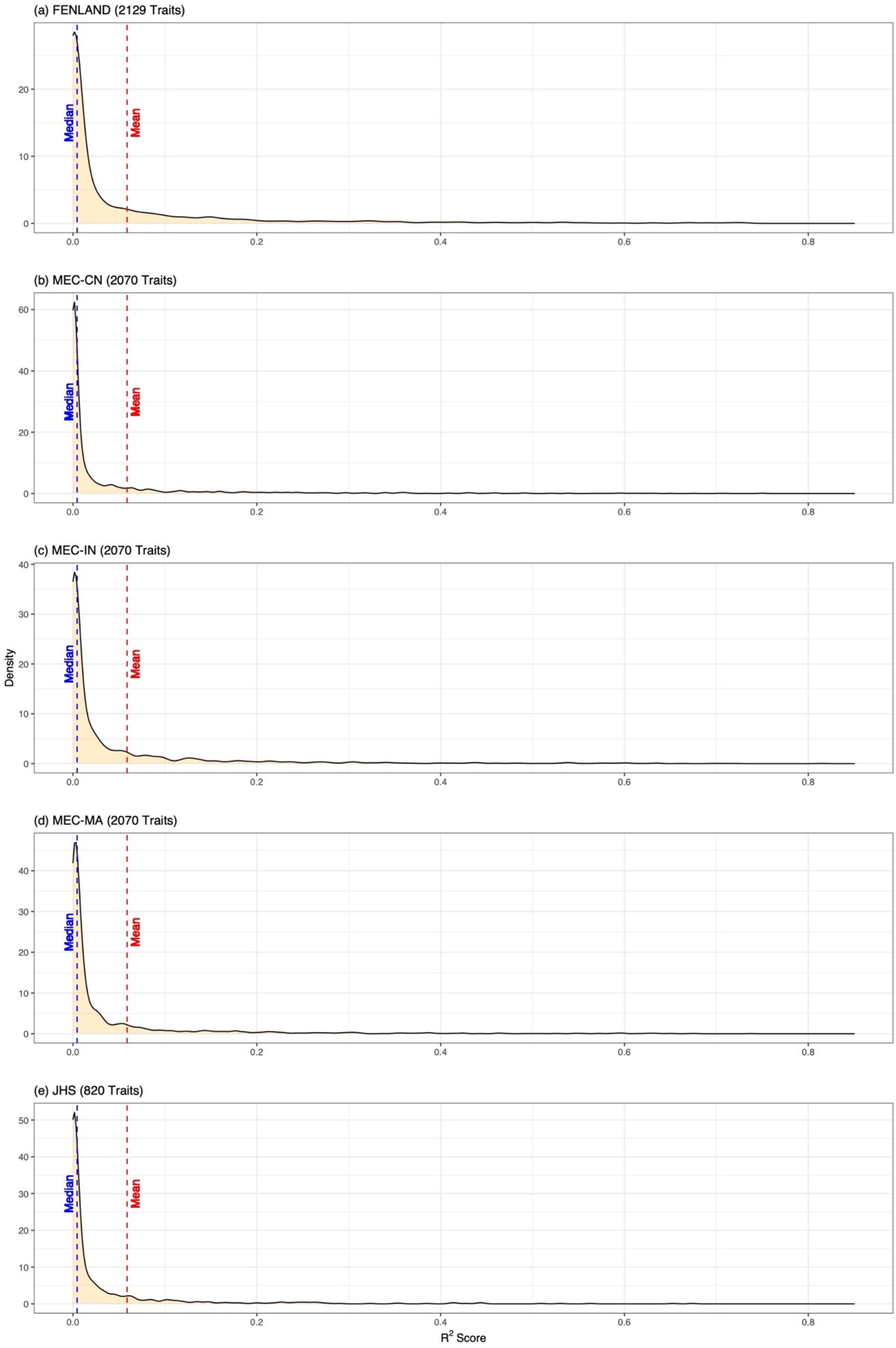

**Figure S10: Distribution of $R^2$ performance in external validation for genetic scores of Olink traits.** This analysis included all the traits validated in the INTERVAL withheld set.

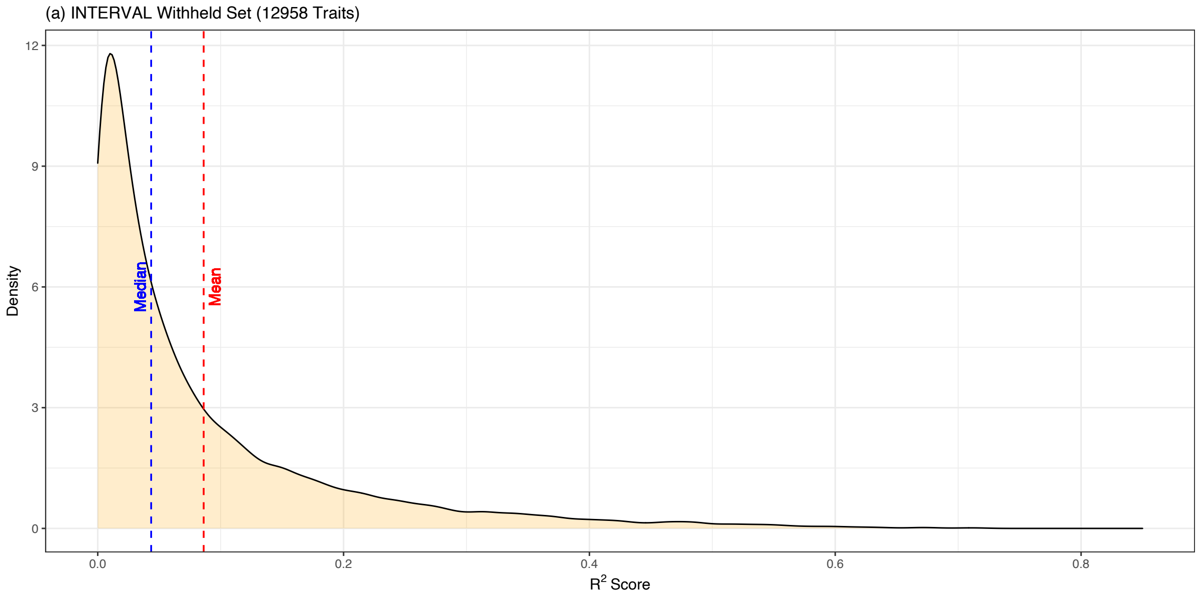

**Figure S11: Validation of genetic scores in external European cohorts.** The figures compare the spearman correlation scores between internal validation and external validation with an European cohort on each platform, in which points are coloured by the variant missingness rate in the external cohort and the blue line shows the linear models fitting the data points.

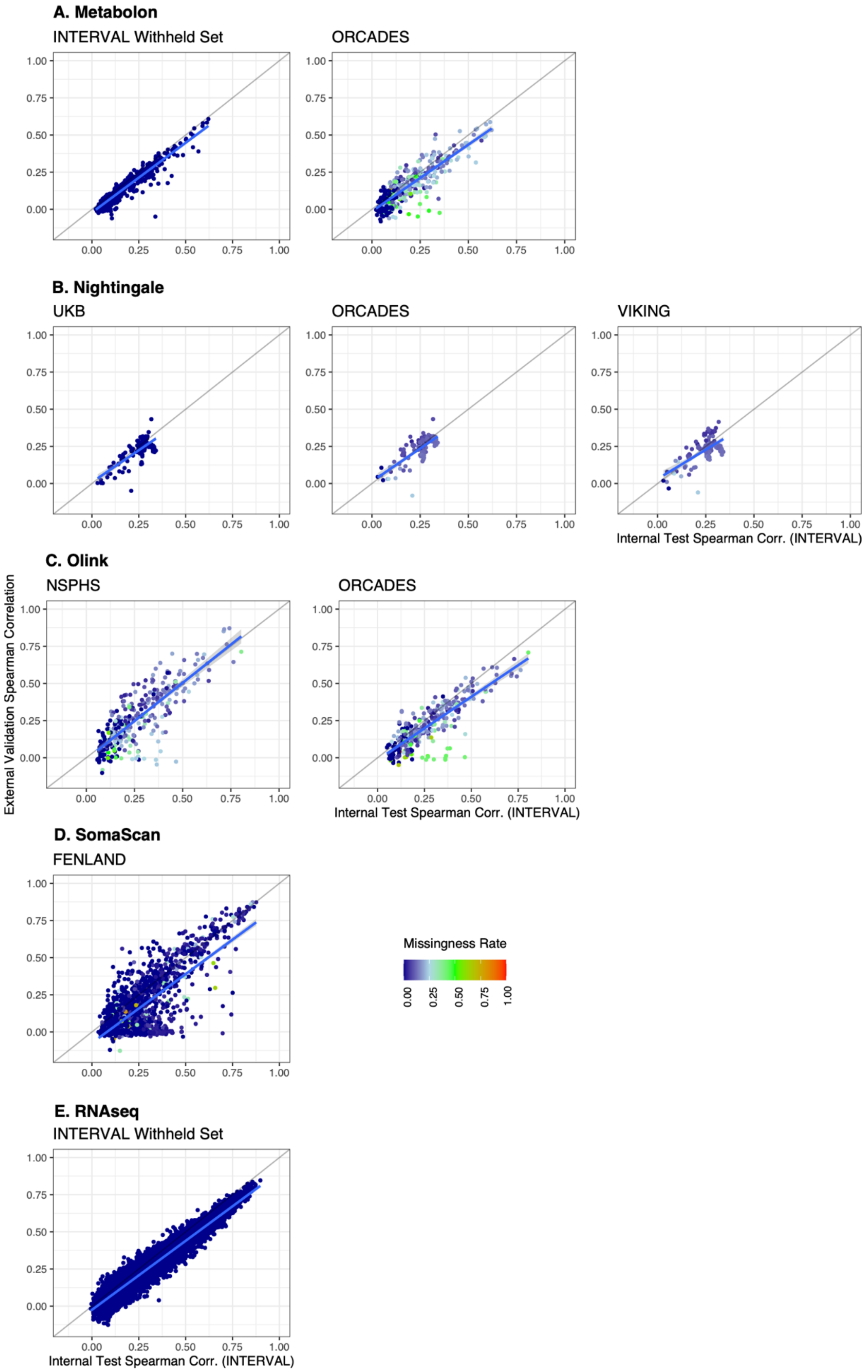

**Figure S12: Validation performance change of genetic scores by their variant missing rates in external cohorts across ancestries.** External validation results in European cohorts were merged in each platform to increase the statistical power in this analysis, which include NSPHS and ORCADES validations for Olink, and ORCADES and VIKINGS validations for Nightingale. Note that INTERVAL withheld subset validations and UKB validation for Nightingale traits were excluded in this analysis due to there is no or nearly no missingness in these validations. Validation results in each platform were ranked by their variant missing rate of genetic score models in the external cohort and grouped into tertiles, where variant missing rate = the number of variants missing in the validation cohort / the total number of variants of the genetic score model. This figure presented the mean and standard error (SE) of $R^2$ performance change of trait genetic scores between internal and external validation across tertiles of validation results. EUR: European; CN: Chinese; IN: Indian; MA: Malay; AF: African American.

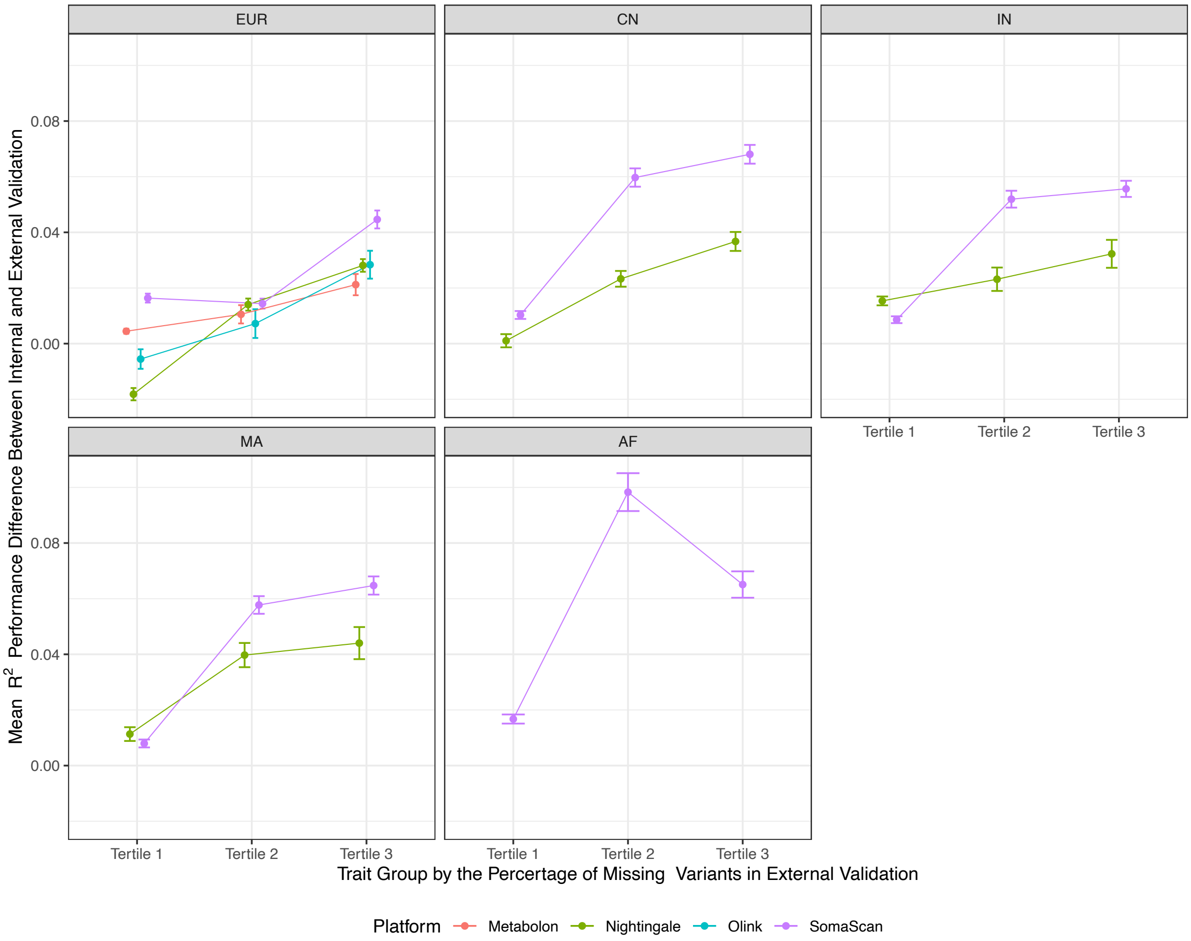

**Figure S13: $R^2$ performance comparison of genetic scores between external European cohorts.** The analyses included all the overlapped traits between two external validations at each platform. The blue line shows the linear models fitting all the performance comparison points.

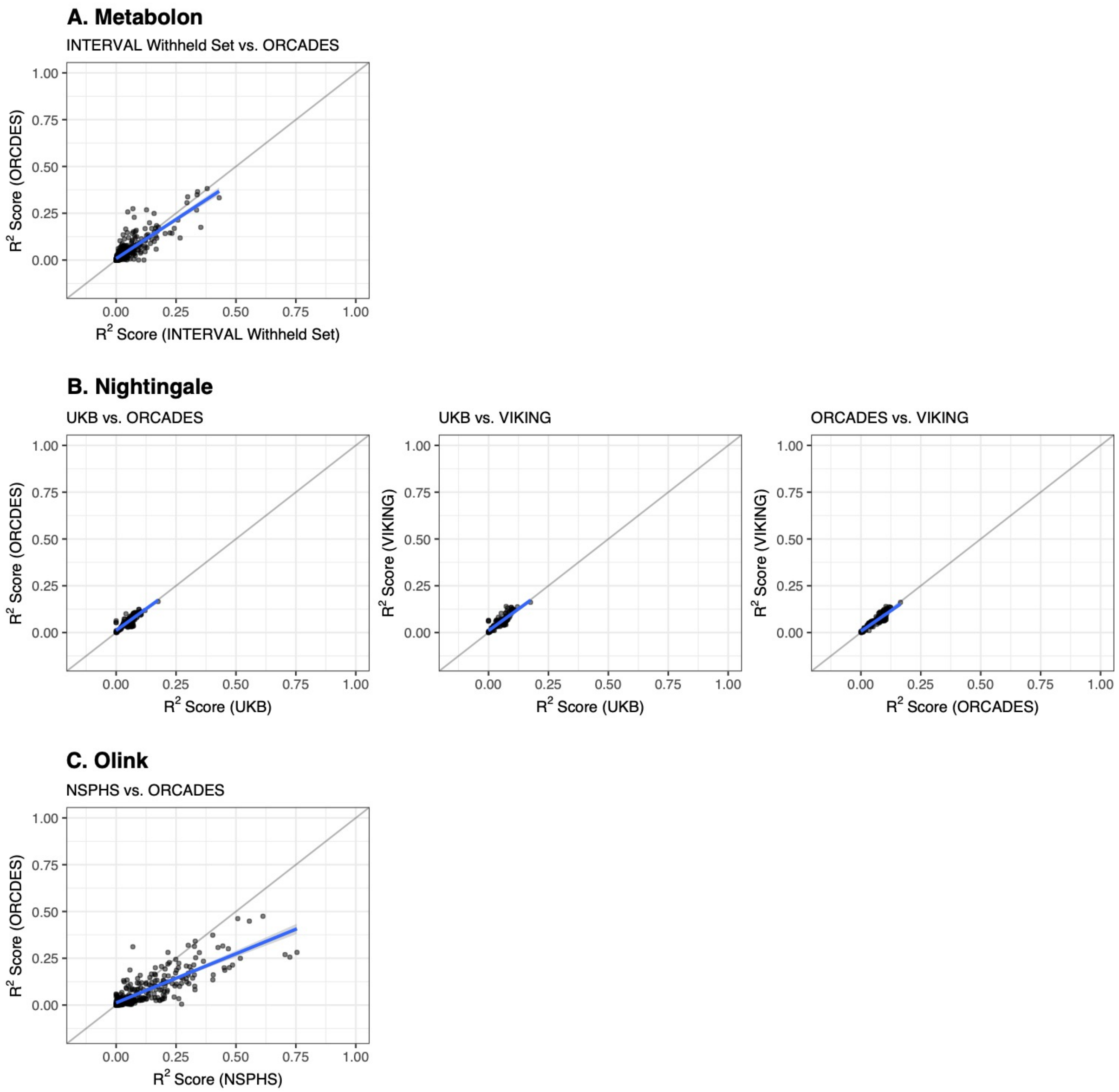

**Figure S14. Key features of online portal for accessing genetic scores of biomolecular traits. (a) Organization of genetic scores on the portal; (b) Example of how biomolecular traits and their genetic score-related information can be explored; (c) Example of how summary statistics of training and validation cohorts are presented; (d) Example of how validation results and genetic score models can be downloaded; (e) Example of how validation results and trait-related information can be visualized.**

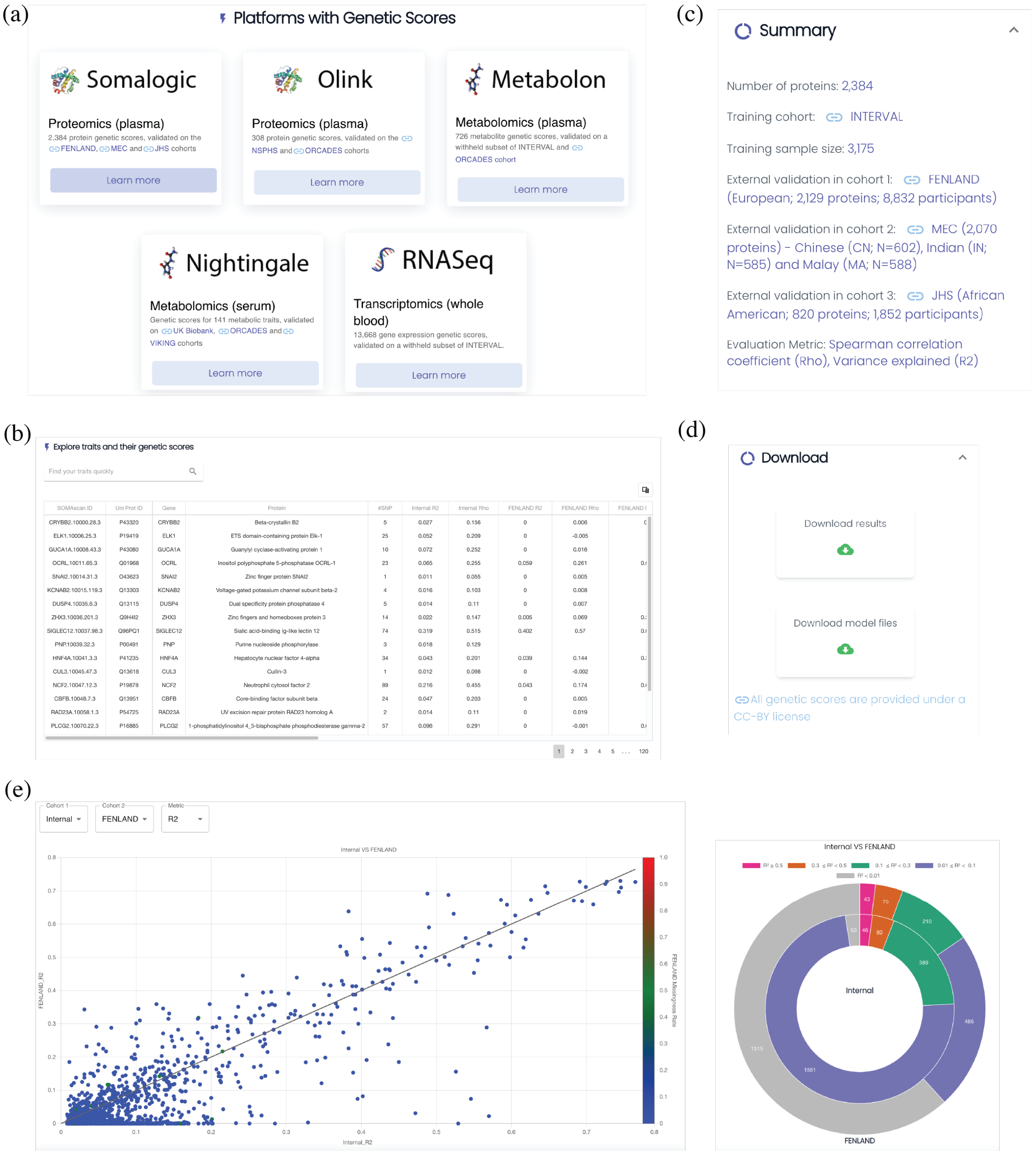

**Figure S15: Coverage analysis for blood proteins in the lowest-level pathways.** This analysis looked at all the lowest-level pathways of super-pathways curated at Reactome. Where at least one protein genetic score are included in the entities of a lowest-level pathway, we considered this pathway is covered by proteins of this study. This figure shows the percentage of the lowest-level pathways a group of proteins (By $R^2$ in internal validation) covered among all the lowest-level pathways of each super-pathway.

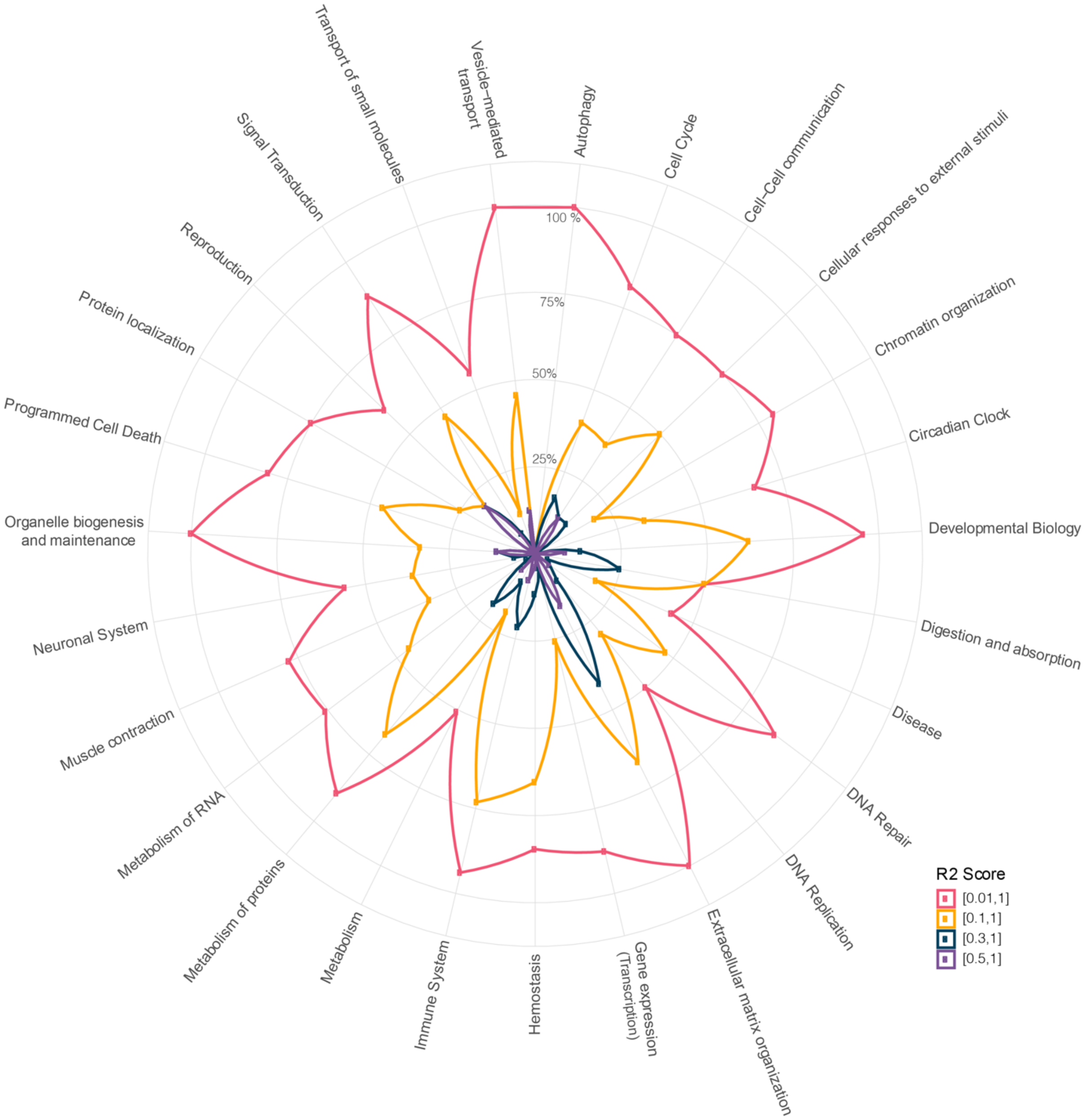

**Figure S16: $R^2$ Performance comparison of genetic scores for shared proteins trained on SomaScan and Olink in INTERVAL.** We compared the internal validation $R^2$ performance of 169 shared proteins on SomaScan (or Olink) with $R^2$ performance of their corresponding genetic scores trained on Olink (or SomaScan) in predicting protein levels on SomaScan (or Olink) using all the INTERVAL training samples. The points are coloured by the Pearson's *r* score between the actual proteins levels of a protein measured by SomaScan and Olink for those samples who were assayed using both platforms.

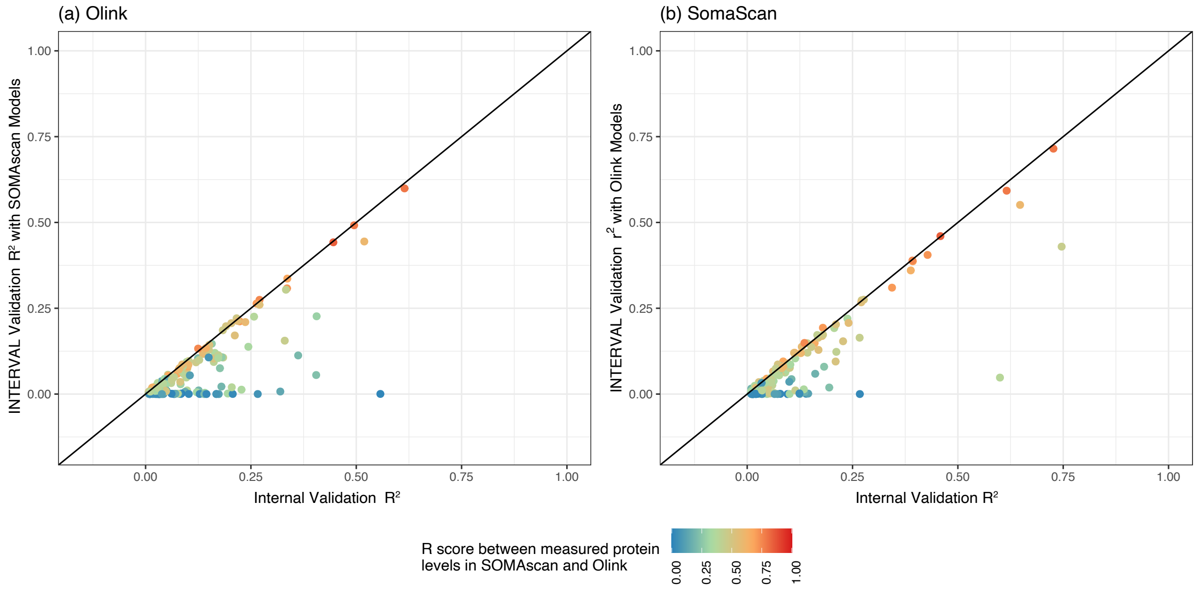
